# Early prediction of declining health in small ruminants with accelerometers and machine learning

**DOI:** 10.1101/2020.08.03.234203

**Authors:** Axel X. Montout, Ranjeet S. Bhamber, Debbie S. Lange, Doreen Z. Ndlovu, Eric R. Morgan, Christos C. Ioannou, Thomas H. Terrill, Jan A. van Wyk, Tilo Burghardt, Andrew W. Dowsey

**Affiliations:** Bristol Veterinary School, University of Bristol, Bristol, UK; Department of Population Health Sciences, Bristol Medical School, University of Bristol, UK; 13 Spey St., Extension 3, North Mead, Benoni 1501, Gauteng Province, South Africa; KZN Department of Agriculture and Rural Affairs, P/B X9059, Pietermaritzburg, 3200, KwaZulu-Natal Province, South Africa; School of Biological Sciences, Queen’s University Belfast, Belfast, UK; School of Biological Sciences, University of Bristol, Bristol, UK; Department of Agricultural Sciences, Fort Valley State University, Fort Valley, Georgia, USA; Department of Veterinary Tropical Diseases, Faculty of Veterinary Science, University of Pretoria, South Africa; Department of Computer Science, University of Bristol, Bristol, UK

**Keywords:** FAMACHA, *Haemonchus contortus*, Anthelmintic Resistance, Precision Livestock Farming, Accelerometers, Machine Learning

## Abstract

Assessment of the health status of individual animals is a key step in the timely and targeted treatment of infections, which is critical in the fight against anthelmintic and antimicrobial resistance. The FAMACHA scoring system has been used successfully to detect anaemia caused by infection with the parasitic nematode *Haemonchus contortus* in small ruminants and is an effective way to identify individuals in need of treatment. However, assessing FAMACHA is labour-intensive and costly as individuals must be manually examined at frequent intervals. Here, we used accelerometers to measure the individual activity of extensively grazing small ruminants (sheep and goats) exposed to natural *Haemonchus contortus* worm infection in southern Africa over long time scales (13+ months). When combined with machine learning, this activity data can predict poorer health (increases in FAMACHA score), as well as those individuals that respond to treatment, all with precision up to 83%. We demonstrate that these classifiers remain robust over time. Interpretation of trained classifiers reveals that poorer health significantly affects the night-time activity levels in the sheep. Our study thus reveals behavioural patterns across two small ruminant species, which low-cost biologgers can exploit to detect subtle changes in animal health and enable timely and targeted intervention. This has real potential to improve economic outcomes and animal welfare as well as limit the use of anthelmintic drugs and diminish pressures on anthelmintic resistance in both commercial and resource-poor communal farming.

---

Livestock farming in resource-poor communities presents multiple challenges such as economic losses from a variety of diseases, including parasitic helminth infections (1). This makes optimal helminth management imperative for a farmer to achieve, but is complex and especially difficult without access to expert help. The gastrointestinal nematode *Haemonchus contortus (H. contortus*) has a particularly heavy impact on small ruminants in tropical and subtropical regions, as these regions provide a favourable environment for the parasite’s development. Each female *H. contortus* produces up to 10,000 eggs per day (2), and these develop into infective larvae in a few days under warm and moist conditions. Re-infection can result in high parasite burdens and acute disease outbreaks often leading to death, especially among young animals (3). The disease is primarily the result of bloodfeeding by adult worms in the gut, leading to anaemia, protein loss, and associated consequences for health, growth and fertility (4). The economic loss due to helminth infection in sheep and goat production is substantial, for example, an estimated $40 million per annum in the Kano area of northern Nigeria and $26 million per annum in Kenya (1).

Although multiple worm control strategies for resources poor farmers exist (5), including chemical dewormers, vaccination, anthelmintic drugs, grazing management, specific diets and ethnoveterinary remedies, they all require high manual labour and other expenses (6, 7). In addition, widespread use of anthelmintic drugs has led to the high prevalence of anthelmintic resistance (AR) in countries such as South Africa (8–10). This is due to farmers relying on anthelmintics as the sole method of control against helminth infection, and poor practices such as treating the entire herd when only a few individuals are affected. Although helminth infections are curable, they are common and exact high ongoing costs relative to other diseases due to the complexity and operational difficulties of effective and sustainable management (11). Indeed, farmers in Sub-Saharan Africa rank helminths as the most important disease in small ruminants, in spite of more visible (*e*.*g*. ectoparasitic) and ostensibly more damaging (*e*.*g*. foot and mouth disease virus) pathogens (12).

A path to more sustainable and efficient control of *Haemonchus* infection is the effective clinical evaluation of individual animals, and selective treatment of those unable to cope (13), leaving the rest untreated. This targeted treatment is preferable because helminths are aggregated among their hosts, such that a few individuals in a group tend to carry the majority of the disease burden, and disproportionally drive onward transmission (14). The untreated individuals provide refugia for anthelmintic-susceptible genotypes, slowing the development of AR (13, 15, 16). The FAMACHA clinical system was developed to deliver on this principle in sheep and goats and consists of a calibrated colour chart against which the colour of the conjunctivae of the eye is compared (17, 18). FAMACHA scores range from 1 to 5, with score 1 being the most healthy, equating to haematocrit *≥* 28%, and score 5 the most severely anaemic (haematocrit *≤* 12%). The scoring system requires minimal training, provides immediate results, and does not rely on expensive equipment or laboratory analysis. However, training is needed, and there is a shortage of qualified trainers, particularly in resource-poor regions. Furthermore, the system relies on frequent close handling and examination of individual animals, which is laborious and costly, given that examination of entire herds is required weekly during high-risk periods (17).

A promising alternative could be to use biologgers to remotely monitor behaviour linked to poor health (19–22). Thanks to advancements in on-animal sensor technology, effective tracking and monitoring of terrestrial animals can now give access to large quantities of data on their behaviour and their interactions with each other and with their environment (23). Accelerometry data has been successfully used to classify livestock behaviour, for example, Moreau *et al*. (24) identified eating, resting and walking in goats by using tri-axial accelerometers. This approach is however sensitive to the sensor position on the animal. Other studies used machine learning approaches. Vazquez *et al*. (25) used a combination of accelerometry and gyroscopic data with online learning to detect sheep behaviour (walking, standing and lying). Scoley *et al*. (26) studied the effect of milk and forage feeding in dairy calves with the IceQube® automatic activity sensors (IceRobotics Ltd., Edinburgh, Scotland, UK). The study revealed that calves displayed activity linked to hunger if they were fed high levels of milk replacement in their early life but were fed reduced levels later. By combining GPS data and 3-axis accelerometry data, González *et al*. (27) were able to accurately detect foraging and travelling behaviour in grazing cattle. Högberg *et al*. (28) measured the accuracy of two commercial sensors (CowScout(GEA Farm Technologies) and the IceTag® (IceRobotics Ltd.) mounted on dairy cows. Both devices use accelerometers to determine lying, standing and walking. The study showed that lying and standing could accurately be detected but walking detection was inaccurate. Although these studies focus on specific activities, all have the potential to be used for livestock management through the monitoring of behaviour. However, by instead using summary statistics derived from limited behavioural classes, for example, the proportion of time walking, much of the rich information in the raw data is lost. In contrast, we use machine learning directly on the full accelerometry data profile for automatically detecting changes in the FAMACHA score of individual animals, with the aim of early detection of declining health (Fig. 1).

**Fig. 1.**
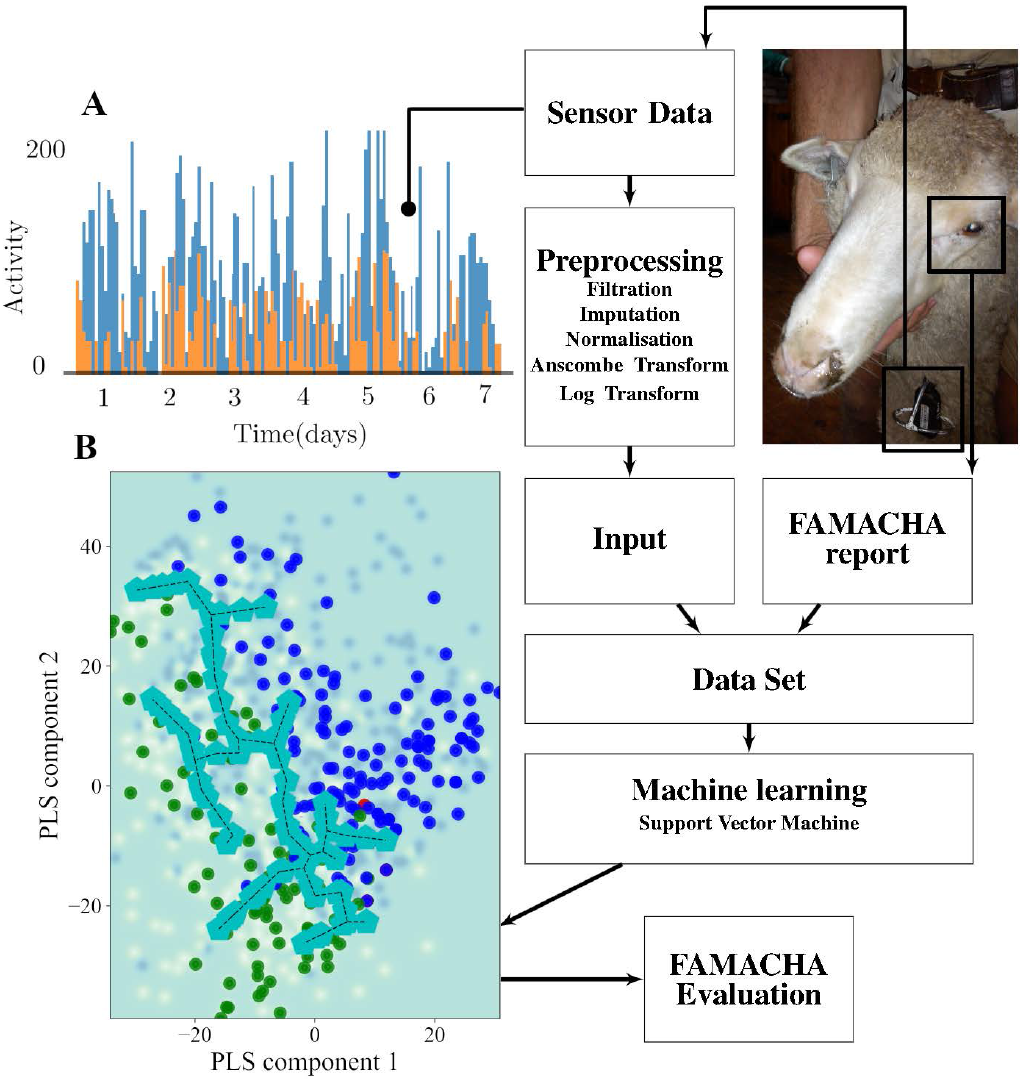
Schematic of our machine learning pipeline. **(A)** The biologger outputs accelerometry data as a count of the instances the acceleration exceeded 2g over a short interval. Here we show derived activity time series for two animals (blue/orange) over 7 days with a bin size of 1 minute. As can be seen, there are significant differences in signal magnitude which we account for through preprocessing. **(B)** The high-dimensional time series data is then combined with the FAMACHA report for supervised machine learning. The scatter plot shows the clustering of the animal samples in this space for training (blue and green points represent healthy and unhealthy animals, respectively). A Support Vector Machine with 5-fold crossvalidation repeated 10 times is then used to derive the classification boundary (turquoise) and to derive predicted probabilities that each sample is healthy or unhealthy.

Our study group consists of a sheep flock in Delmas, Mpumalanga Province, South Africa, and a goat herd at Cedara Government Agricultural Animal Production Research Farm, Howick, KwaZulu-Natal, South Africa (Table 1). A low-cost transponder containing an accelerometer was suspended by a sturdy ribbon around the neck of each animal and used to transmit activity levels in real-time to a base-station continuously for more than a year (Table 1, Fig. S1). During this time, the FAMACHA score was assessed every week for sheep, and every two weeks for goats. A supervised machine learning pipeline with deep-learning based imputation was developed to predict an increase in FAMACHA score from 1 (“optimal”) to 2 (“acceptable”), as well as from 2 to 1 after individual anthelmintic treatment (Fig. 1) . Classification drift including robustness to seasonality was assessed through temporal validation. We also performed inverse transformation of the trained classifiers for visual interpretation of the behavioural cues that distinguish healthy animals from those with a trajectory to poorer health.

**Table 1.** Characteristics of the study group.

|  | Cedara | Delmas |
| --- | --- | --- |
| Data collection period | Mar 2015 to Apr 2016 | Apr 2012 to Jul 2013 |
| Species | Sheep | Goat |
| Animals tagged | 64 | 227 |
| Tag type | Accitrack v2 | Accitrack v1 |
| Age range | 2-6 years | 2-6 years |
| Average weight | 72.79 kg | 44.19 kg |
| FAMACHA evaluation | weekly | fortnightly |
| Animals evaluated | 13 | 32 |
| FAMACHA 1 → 1 | (33.4%) 414 | (20%) 79 |
| FAMACHA 1 → 2 | (19.2%) 238 | (11.6%) 46 |
| FAMACHA 1 → 3+ | 0 | (5.0%) 2 |
| FAMACHA 2 → 1 | (18.4%) 229 | (13.0%) 50 |
| FAMACHA 2 → 2 | (28.6%) 355 | (19.9%) 79 |
| FAMACHA 2 → 3+ | 0 | (10.3%) 43 |
| FAMATHA other | 0 | (20.2%) 80 |

## Results

### Classifying health status

#### Repeated K-Fold cross validation

We first aimed at predicting which individuals were in persistently poorer health, i.e. have a FAMACHA score of 2 *→* 2, based solely on the 7 days of accelerometry data immediately prior to the state ( 2 *→* 2). For this we had 142 cases of FAMACHA 1 *→* 1 and 140 examples of FAMACHA 2 *→* 2 for the sheep at Delmas. With these annotations, we trained and tested our supervised machine learning pipeline using 5-fold cross-validation repeated 10 times to provide estimates of model performance with low bias and variance. The machine learning pipeline was able to predict the presence of FAMACHA score 2 *→* 2 with median precision of 74% (75% and 73% for the healthy class and the unhealthy class, respectively). In a Receiver Operating Characteristic (ROC) analysis, the median area under the curve (AUC) was 83% (Fig. ).

For the goats at Cedara we had 79 examples of FAMACHA 1 *→* 1 and 77 of FAMACHA 2 *→* 2. The median precision was 63% (64% and 62% for the healthy class and the unhealthy class respectively) for individual goats declining in health. In the ROC analysis, the median AUC was 70% (Fig. S28). The observed lower prediction power is likely due to the smaller number of examples for the goats.

#### Leave two out cross-validation at the animal level

To test the model for prediction on completely unseen animals, we proposed a leavetwo-out (LTO) cross-validation approach. We chose to exclude 2 animals instead of 1 due to the small number of individuals (only 13) which would have limited the number of folds available to assess model uncertainty. For the sheep farm where the dataset was larger, we trained a linear SVM model with 7 days of accelerometry data from all the sheep data available excluding each combination of 2 individuals, which were held out to validate the model. Similarly to the K-fold approach, we aimed at predicting which individuals would have a FAMACHA score of 2 *→* 2 or 1 *→* 1; for this, we had respectively 127 and 126 examples.

The machine learning pipeline was able to predict the presence of FAMACHA scores 2 *→* 2 with a median precision of 63.5% (64% and 63% for the healthy class and the unhealthy class, respectively). In a Receiver Operating Characteristic (ROC) analysis, the median area under the curve (AUC) was 71% (Fig. ).

#### Classifying the response to treatment

As part of routine husbandry, each animal which scored 2 or more during a FAMACHA evaluation was immediately treated with Levamisole (Ripercol-L, Bayer Animal Health) at 7.5 *mg*.*kg*^*−*1^ (29). We created a dataset to examine our ability to predict which animals responded well to treatment, comparing FAMACHA 2 *→* 1 against FAMACHA 1 *→* 1. This was based on classifying the 7 days of accelerometer data immediately following treatment for the sheep. For the goats, FAMACHA 2 *→* 1 against FAMACHA 1 *→* 1 was also compared and 2 day of accelerometer data was used.

For this analysis, we had 90 examples of FAMACHA 2 *→* 1, 142 for FAMACHA 1 *→* 1 for the sheep, and 53 examples of FAMACHA 2 *→* 1 and 79 examples of FAMACHA 1 *→* 1 for the goats.

This modest training set size may diminish the performance of the machine learning (Fig. S22. FAMACHA transitions available). Nevertheless, the classifier was able to predict a drop in FAMACHA score indicating a response to treatment with median precision of 53% and 55% for the sheep and goats, respectively, and no change in FAMACHA score with median precision of 70% and 71%. As shown in (Fig. E, F and Fig. S29, the median AUC was 66% for the sheep and 71% for the goats.

#### Temporal validation to assess concept drift

The notion of “concept drift” describes the decrease in performance of a given classifier due to changing environmental or sensing conditions over a period of time. For example, training data collected at the start of a given period becomes less representative of future data. This is a common issue in long-duration supervised classification problems that use real-life data which is, in most scenarios, changing intrinsically (25). An analysis was devised to test how much concept drift affects our findings. Two equally-sized periods of data were extracted from the sheep and goat datasets to maximise seasonal differences: for the sheep farm,(i) the first period from March 2015 to October 2015; (ii) the subsequent period from October 2015 to April 2016. The first period contained 287 (150 healthy and 137 unhealthy) samples using 6 days of activity data and 287 samples (148 healthy and 139 unhealthy) in the second period. For the Sheep farm, it is clear from observing the resulting scatter plots of the trained classifiers (Fig. G, H for the second period, Fig. S30 for the second) that while there is little observable drift in the activity of the healthy animals, the less healthy animals cluster differently depending on time period. Nevertheless, these clusters do not interfere with the decision boundary and hence the overall precision of prediction did not noticeably decrease, yielding a median AUC of 69% for the sheep when training on the first period and testing on the second, and 75% when doing the reverse.

For the goat farm, we had only 46 (26 healthy and 20 unhealthy) samples for the first period and 51 (14 healthy and 37 unhealthy) samples for the second one. Because of the limited number of samples, latter results are inconclusive (see Fig. S25).

#### Adding rainfall

*H. contortus* is well known to hatch after humid, hot weather, and to require rainfall for movement onto pasture (30, 31). We therefore tried training models on humidity, temperature, windspeed and rainfall, determining that rainfall alone was optimal (Fig. S27) and hence validated models using rainfall only, activity only, and with both rainfall and activity. We used the Wilcoxon signed-rank test to compare the per-fold AUCs between these models (Fig. S26).

For the Sheep farm, results on activity covariates alone showed statistically significant improved performance than with rainfall alone (p*≤*0.001), and also improved performance over considering both activity and rainfall together (p=0.025) (Fig. 3C). For the Goat farm, results on activity covariates alone showed statistically significant improved performance (p=0.013) compared to rainfall alone, with no evidence that performance differed compared to using activity and rainfall together (Fig. S25). We hypothesise that when there is a high enough sample count for accelerometry data such as in the sheep farm, the information provided by the rainfall data is already intrinsically contained within the activity data. For the sheep farm, we had 75% AUC with rainfall only and 81% for activity and rainfall, while on the goat farm, we had 68% and 70% respectively.

#### Interpreting the classifier

In order to understand how our model discriminates between animals on healthy and less healthy trajectories, we analysed the output of the classifiers in Fig. 2 B by multiplying the derived SVM feature weights (32) which define the classification boundary. As illustrated in Fig. 3B, the result shows that the trained model ascribes more importance to activity during the night for the sheep farm (Wilcoxon p=0.015). For the goat farm, the results were inconclusive (Fig. S39).

**Fig. 2.**
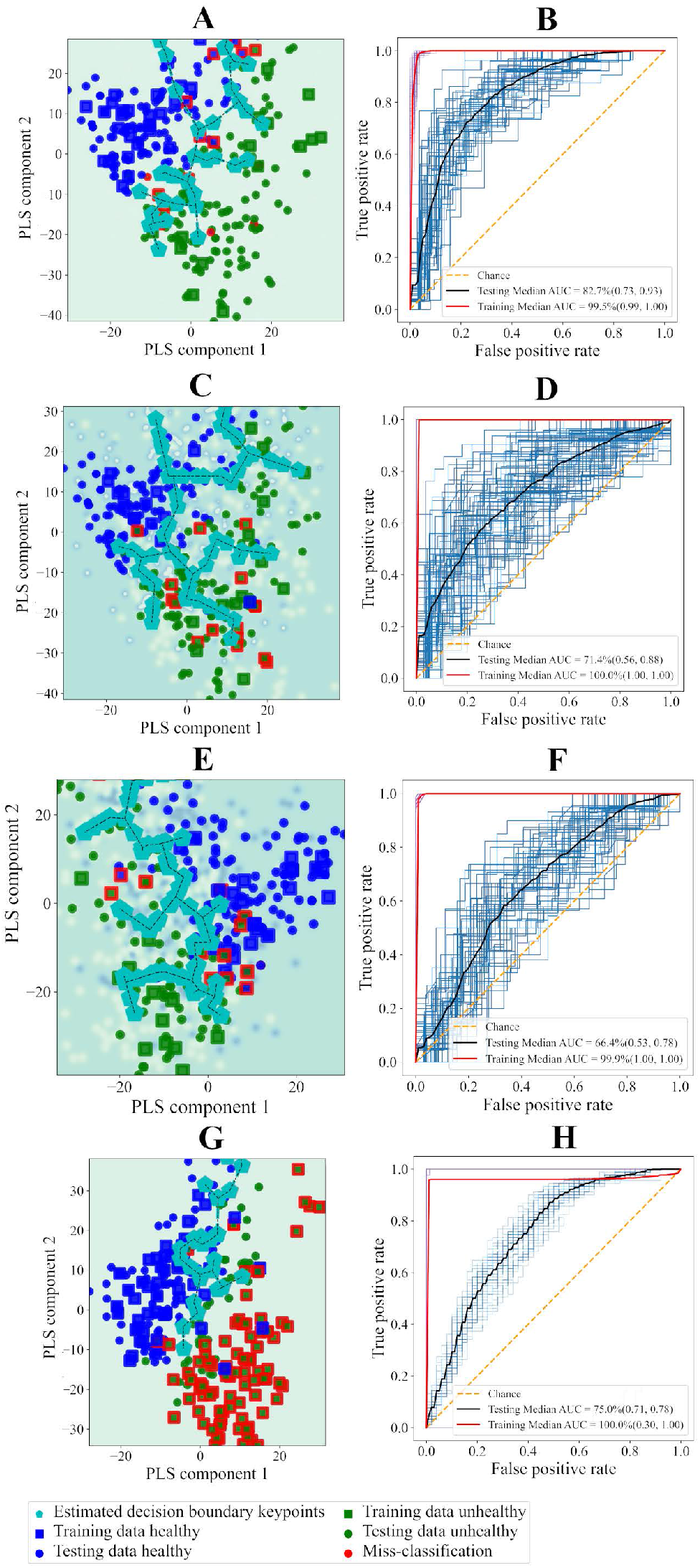
Decision boundary plots and ROC curves for the Sheep farm. (A) and (B) shows the result of the health status experiment with K-Fold cross validation, (C) and (D) for Leave two out cross validation. (E) and (F) show the response to treatment while (G) and (H) show the temporal validation for training on the second period and testing on the first.

**Fig. 3.**
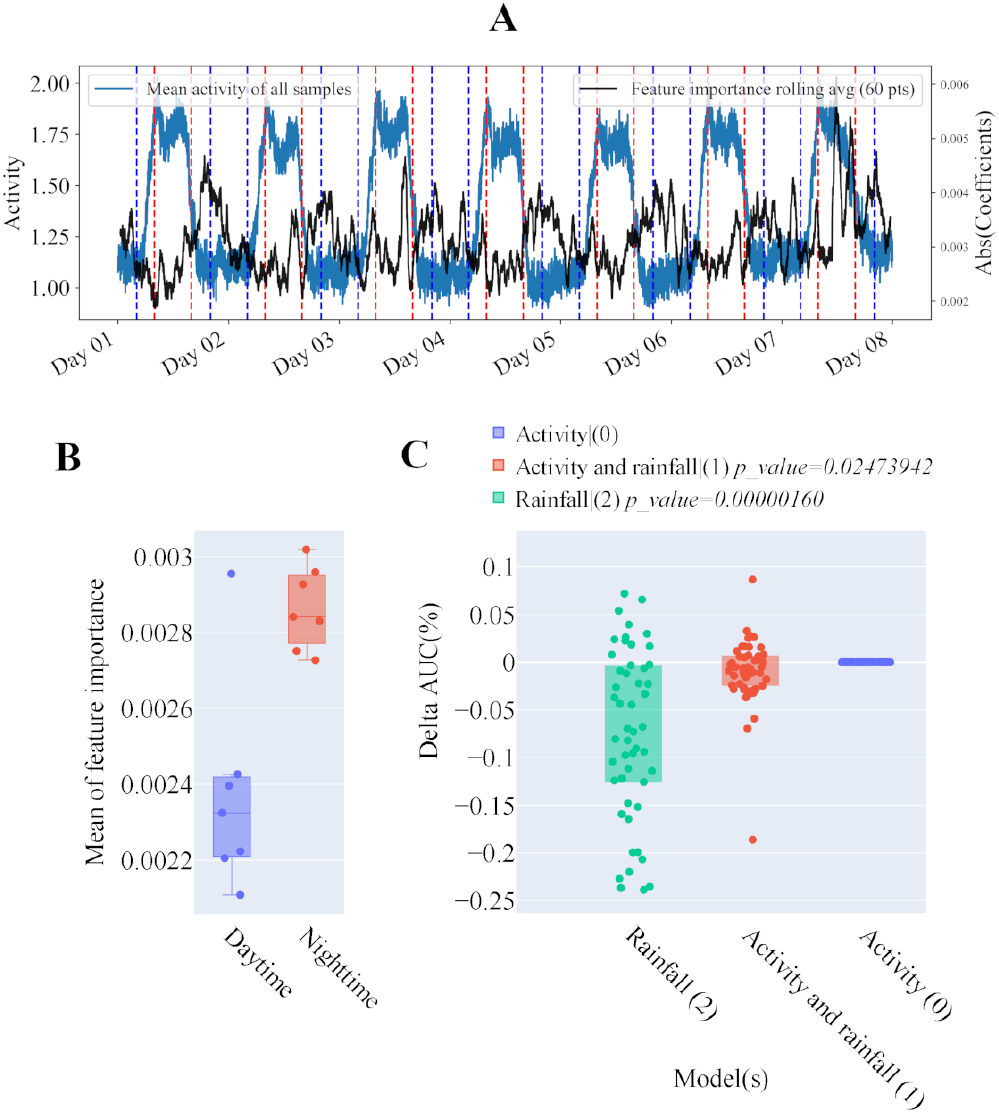
Interpretation of the classifiers from Fig. 2 A,B. After fitting the SVM model (A) with the 7 days post-proccessed time series, the blue trace shows the mean of all the samples. We can display the average of the feature importance attributed by the classifier, “black trace”. **Statistical analysis** (B) Illustrates the mean importance of all 7 daytime periods vs all 7 night time periods, daytime being defined between the latest sunrise and the earliest sunset, 8am to 4pm, illustrated by the vertical dotted blue and red lines respectively. **Performance of SVM model trained on activity against Rainfall or both**. (C) The left y-axis shows the number of samples while the right axis shows the difference compared to the best model AUC.

#### Discussion

Our analyses reveal that an increase in FAMACHA score from 1 (“optimal”) to 2 (“acceptable”) and remaining at 2 for up to one week for sheep and two weeks for goats, which is considered sub-clinical disease, can be predicted from behaviour measured using low-cost biologgers and that this prediction is improved over using climate data alone. We discovered that the discriminative ability of our classifier increases during the night for the sheep, with inconclusive results for the goats possibly due to the smaller sample size.

While efforts in developed countries have been focused on building high-precision, securely mounted and precisely fitted sensors, we demonstrate that robust results can be gained from much simpler, low-cost systems with rudimentary maintenance requirements suitable for both commercial and RP farmers in developing countries. It is important to note that due to significant calibration and mounting variation between transponders, including loosening of the transponder over time, it was necessary to perform normalisation of each activity trace to the herd/flock mean. This meant that uniform reductions in activity level from week to week are likely to be normalised out of our data. Nevertheless, a biological reason for a completely uniform reduction in activity level is implausible; instead, the intensities of some daily activities are likely to be impacted more than others. We have shown that changes in the variation of activity levels constitute a strong predictor of early changes in health status, as regards haemonchosis and are robust to technical variation.

In this study we have focused on accelerometry data and climate data, but other exogenous factors such as body weight etc could be beneficial to the prediction. Nevertheless, optimal incorporation of these data types is challenging because of their potential non-linear and/or lagged or cumulative effect on health. Conversely, the fundamental advantage of high-dimensional longitudinal data from accelerometers is that the end effect of these covariates could intrinsically be contained within the data directly, which machine learning approaches have the potential to deconvolute. The ability of machine learning to detect health-relevant changes in behaviour under variable climatic conditions could make it especially useful as climate change drives increasingly unpredictable transmission patterns among helminths (33), and hence to support adaptation to climate change by resource-poor farmers (34).

Collecting robust annotated datasets is especially challenging in resource-poor farming systems where farming practices are generally less consistent, regulated and well-funded. Because of this and the need for intensive manual labour over a prolonged period, our datasets are highly valuable. Although the training data obtained is dependent on farm topology, location and management, we have shown that a basic machine learning pipeline can discriminate on behavioural cues dominated by fluctuations in activity levels. Some concept drift was found, particularly among animals with increasing parasitic burden. This suggests two broad avenues for future research: (a) Starting from deployments of our pre-trained model, use of online reinforcement learning techniques to create a ‘life-long learning’ decision support system which identifies animals for FAMACHA evaluation and feeds back the results to dynamically update the model calibration and improve future predictions; (b) Multivariate time-course statistical modelling to further characterise the nature of sheep and goat behaviour in health and disease.

Notably, in this study, we have focused on FAMACHA evaluation of *H. contortus* infection; whether the multi-label classification of a range of different disease states and transient events is possible is currently unknown but would require extensive studies to attain the appropriate predictive power. In addition, as our accelerometry data is based on a simple activity count paradigm, we hypothesise that activity traces could be derived from other sensor types, such as video, for direct input into our prediction model.

Helminths negatively impact livestock productivity worldwide, and in resource-poor settings are considered ‘neglected cold spot’ diseases, in that they are preventable in principle, but farmers continue to struggle to manage their effects (35). Technical improvements in helminth control consequently have especially high potential to positively impact farmer livelihoods, with knock-on benefits for human nutrition and health (36). The FAMACHA system has been successfully adopted by smallholder farmers in Africa, but sustained use is difficult because of high training and labour requirements (37). Our results show that it is feasible to apply machine learning approaches to data streams that are attainable on smallholder farms in Africa to detect early changes in health status and support timely and targeted intervention.

## Methods

### Study group

Data were collected from a farm of 108 acres close to Delmas in Mpumalanga Province, South Africa, and at Cedara, a government research farm and agricultural college, the pastures of which comprise of 25 acres adjacent to Hilton, KwaZulu-Natal. As described in Table 1, 31 female adult *Ile de France* sheep ewes at Delmas and 64 goats at Cedara were individually FAMACHAevaluated (17) at weekly or fortnightly intervals, respectively, and had associated longitudinal accelerometry data recorded within the study period. On both farms, there were multiple improved pastures, which were irrigated, that were used alternatively at intervals according to visual assessment of amounts of available water and forage. At both farms, young adult ewes/does were randomly selected for the trials, without attention to reproductive class, but remained with their flocks/herds of origin for the duration of each trial. The animals were kraaled at night and let out at a standard time in the mornings for herding to pasture, where they remained until collected and returned to the kraals in late afternoon. Adjustments were made according to season and special management events such as vaccination and hoof inspection, which were conducted first thing in the morning. As part of routine husbandry, each animal which scored *≥* 2 during a FAMACHA evaluation was immediately treated with Levamisole (Ripercol-L, Bayer Animal Health) at 7.5 *mg*.*kg*^*−*1^ (29).

### Telemetric monitoring

Telemetric monitoring systems were supplied by Accitrack Ltd., Paarl, South Africa. Tagged animals on both farms were equipped with low-cost RFID transponders suspended by a sturdy ribbon around the neck (Fig. 1). A single solar-powered base station was installed on each farm, mounted at the top of a five-meter wooden pole (Fig. S1). Each transponder contains an active RFID transceiver operating at 868Mhz as well as a battery and an A1-type accelerometer for measuring activity level. The accelerometers had a set acceleration threshold of 2g so that every time an acceleration *≥* 2g is sensed, a stored integer is incremented by +1. Version 2 transponders had an increased range (10km versus 1km) and a larger battery, at the expense of significantly increased weight. All tags were set to transmit the data every minute to the base station, at which time the accelerometer count was reset to zero. In order to extend battery life, the tag only transmits data to the base station once a minute, with data transmitted including the identifier of the transponder, the battery level, the signal strength, a timestamp, and the activity level. Data transmission is not performed if the signal-to-noise ratio drops below 10dB, which can occur when there are significant occlusions between the animal and the base station. In these cases, the data for that time interval is lost. Through mobile connectivity with General Package Radio Services (GPRS), the base station then regularly forwards the received data to the Accitrack cloud repository.

### Data management and visualisation

All raw data is stored on the cloud repository for two weeks only due to storage limitations, hence it was manually downloaded regularly by a researcher at the University of Pretoria for archival (29). The exported data took the form of Excel spreadsheets containing the sensor outputs in the desired time frame. In cases where the data was not retrieved from the cloud infrastructure, datasets for that time period were lost. For this work, we parsed the raw Excel data into an SQL database. The table storing the raw data recorded at the original minute resolution contains 40,659,086 records. The data was rebinned into multiple time resolutions (Δ*t*) for efficient interactive visualisation. This tool was developed for exploratory analysis to allow us to determine whether transponders were faulty and failed to transmit and provided visual verification of expected behaviours such as decreased activity levels at night. The visualisation also revealed the need for data pre-processing, as changes in average activity signal amplitude across the herd were observed due to varying calibrations of the sensors. In addition, the mounted position of the sensor on each animal influences the sensor’s measurement of acceleration, *e*.*g*. a looser collar would allow broader movement of the sensor and thus higher activity values. In contrast, long wool tended to reduce the number of occasions on which activity was registered.

The data processing pipeline including pre-processing, imputation and machine learning are summarised below, with a detailed description in the supplementary material.

### Pre-processing

We consider a set of animals *r* = 1, 2, … *n* with associated FAMACHA scores *f*_*r,d*_ evaluated on days *d* = 1, 2, … *T*, where *T* is the total number of consecutive days being sampled. Each FAMACHA evaluation is associated with a trace of activity counts *a*_*r,d*_(*t*_*i*_) over time intervals *t*_*i*_ where *i* = 0, 1, … *τ*, where *τ* is the total number of samples within a day (*d* with a temporal resolution of Δ*t*) either preceding or proceeding the evaluation depending on the analysis. Only traces containing a low percentage of missing data and zeros (thus indicating the collar was attached to the animal and functioning correctly) were retained. If the traces exhibited either 50% of the values to be zeros or 20% of the values to be nonexistent, then these were dismissed.

To correct for scaling differences due to device calibration and mounting differences between animals, and the mounting loosening over time, we first define a mean “herd-level” activity trace across all animals with FAMACHA evaluations on the same day:

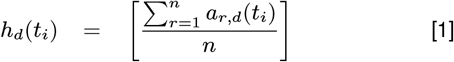

A scaling coefficient for each trace *s*_*r,d*_ is then derived as the median of the trace after having divided it by the herd-level trace:

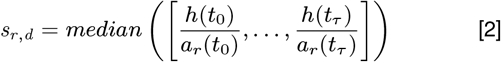

Normalised activity traces *b*_*r*_ are then derived by multiplying each original trace by its respective scaling coefficient:

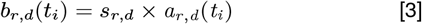

Because our activity data are counts and hence more closely follow Poisson statistics than the Gaussian distribution expected by conventional machine learning methodology, after normalisation we perform an Anscombe transform (38) for variance stabilisation to an approximate Gaussian distribution:

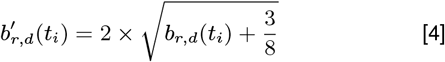

Imputation

Imputing the data saved by an individual transponder without considering the data saved by the other transponders in the herd is intrinsically limiting the ability of any algorithm to accurately predict the missing data points as only the data from this transponder is used. In our study, we can take advantage of the deep-learning based Multi-Directional Recurrent Neural Networks (MRNN) imputation technique devised by Yoon et al (39) which takes advantage of the temporal component in multiple data streams. In our case, this approach can take advantage of the information contained in all of the transponders in the herd to impute the missing data.

It is important to train our imputation models with high-quality data that reflects best the real-world activity of the animals. For this research, the data collection not only suffered from missing data points but also invalid and weak signals (Fig. S3) due to the transponders not being used correctly or placed on the animals securely. For each transponder, we evaluate the complexity (40) of the accelerometry data with the Shannon entropy (41), to select the transponder that contains the densest information and dismiss the lower quality (i.e, a large amount of missingness and abnormal data points) traces. The Shannon Entropy can help us measure the information content in the transponder which ultimately allows us to detect invalid transponders. The two main problems the transponder traces suffer from are the repetition of the same activity count value for an extensive period of time and the excessive amount of missing points. This approach is effective at detecting both.

Fig. S21 highlights how well MRNN learned the data structure of each transponder as it can impute entire sections of missing herd data, for example for the 7th day or at the end of the x-axis. We describe a detailed investigation compared three data imputation techniques in the supplementary material.

### Machine learning

The scale of labelled data for classification was not great enough for deep learning - we therefore investigated a number of classical approaches, with the Support Vector Machine (SVM) (42) demonstrating best performance (see supplementary material). Training/testing datasets for this supervised machine learning technique were constructed using the FAMACHA scores and the standardized activity traces. Repeated nested *k* - fold cross-validation was used to optimize the hyperparameters and evaluate the model. Kim *et al*. (43) showed that the repeated cross-validation estimator outperforms the non-repeated version by reducing the variability of the estimator and providing lower bias. Hence we choose to use 10-times repeated 5-fold crossvalidation to assess a realistic estimate of the performance of the model predictions while making the most of the dataset available. For regularisation, we used a grid search approach to find the optimal SVM parameters ( refer to supporting appendix S6 “Regularisation” for further methodology description). Each ML experiment (classifying health status, the response to treatment and temporal validation) was individually optimised, with the optimal dataset length and models. In the Temporal validation experiment (Fig. S30) to evaluate the models instead of using a simple test/train split we randomly sampled 90% of the samples for training and used the remaining 10% for testing and repeated the process 50 times, allowing us to create an uncertainty range. A detailed investigation into the machine learning pipeline is given in the supplementary material, including justification for the use of SVM and the pre-processing methods described above.

### Supporting Information Appendix

Source code and the datasets are available from https://github.com/biospi/PredictionOfDHealthInSR

## Acknowledgments

We would like to thank the farmers who so diligently assisted with data collection, and Rachel Coetzee and Jan van Rensburg for trial and data management and support. This work was supported by UK Research and Innovation BBSRC grants BB/S014748/1 and BB/H00940X/1, and The Alan Turing Institute under EPSRC grant EP/N510129/1. We also acknowledge support through research grants from Red Meat Research Development South Africa (RMRDSA) and the John Oldacre Foundation through the John Oldacre Centre for Sustainability and Welfare in Dairy Production, Bristol Veterinary School. The Fulbright Specialist Program supported THT on an assignment in South Africa related to this work.

## Notes

https://github.com/biospi/PredictionOfDHealthInSR/blob/master/PNAS_Template_for_Supplementary_Information_2024_May.pdf

### Competing Interest Statement

The authors have declared no competing interest.

### Summary of Updates

We confirmed our findings with the additional cross validation of our model by using leave two out cross validation, a version of leave one out cross validation which allows us to test our model on completely unseen animals. Furthermore, we also show that although the use of exogenous factors such as rainfall has some level of predictive power in our model, using accelerometry data yield better results.

