## Supplementary material for "Early prediction of declining health in small ruminants with accelerometers and machine learning"

#### This PDF file includes:

Supporting text
Figs. S1 to S39
Tables S1 to S4
SI References

### Supporting Information Text

### 1. Visualiation and pre-processing of the raw data

**A** – Sheep wearing a transponder **B** – Base station

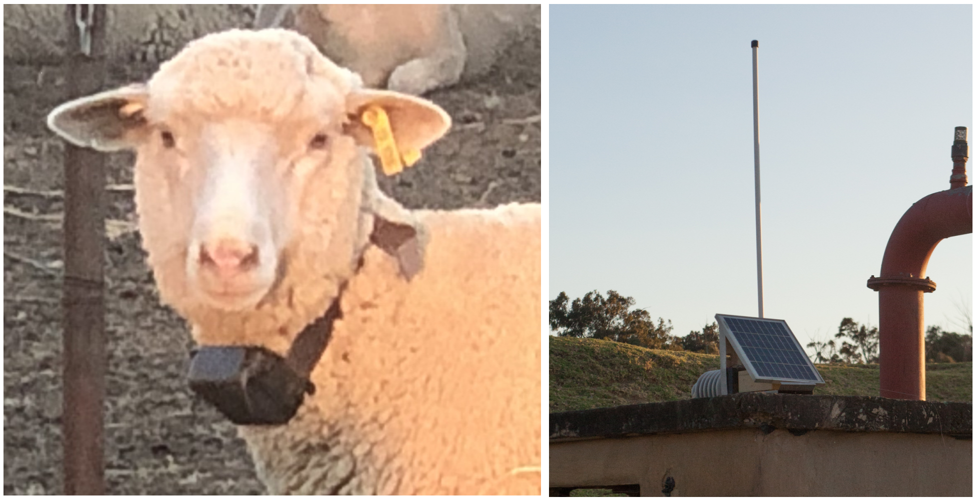

**Fig. S1. Telemetry hardware**. While (A) shows one of the collar mounted v2-accelerometer based transponders used in this research, (B) show a solar panel base station on the roof of a small building on a private South African farm.

**A. Misalignment.** Because of apparent misalignment in the accelerometry record timestamp of the Goat and Sheep transponders, the raw data needs to be "repaired" before imputation. The sampling resolution of the sensors used for this study was set to 1 record per minute. However, the transmission of the data does not always follow the same pattern (1 activity record sent every minute) because of communication errors the sensor transmits data only when possible. Furthermore, it is possible for the sensor to send multiple records for the same minute 'to catch up'. The reason why this occurs is unknown. Because of the irregularities of the sensor output, the first preprocessing step applied to the raw data is to re-align the sampling resolution to 1 record per minute, i.e allocate one record to a single minute bin. The repair process starts by fixing the time axis of every transponder on a common time axis that start at the time of the earliest record across all transponders and similarly ends at the time of the last record. The new time axis will holds a single activity count per bin (Fig. S2).

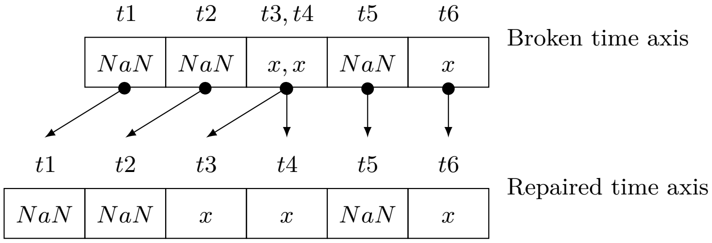


**Fig. S2. Illustration of the back-fill repair process**. In this schematic $x$ is an integer corresponding to a activity count value while 'NaN' (not a number) stands for a missing count.

**B. Transformations.** In general, ML algorithms and statistical models assume that the data have Gaussian residuals. Because these algorithms are sensitive to the input data distribution, it is important to understand the distribution of the data before imputation or classification to optimise our preprocessing approach. Accelerometry data that has been captured as counts naturally follow a statistical Poisson distribution (1) which is a discrete distribution i.e. only takes a discrete set of values. It describes the number of events in a fixed time interval, for example, the number of sales a merchant makes every hour follows a Poisson distribution. This distribution is also characterised by the expected number of events per time interval parameter $\lambda$, where crucially the variance rises directly with the expected number of counts. In addition, it is bounded by 0 and $\infty$. The Poisson distribution assumes that the rate at which each event occur is constant, in other words, the probability of an event occurring in a certain time interval should be the same for every time interval of equal length (assuming the underlying rate does not change due to covariates). And the occurrence of one event does not affect the occurrence of a subsequent event, i.e.

the events are independent. The probability mass function $P$ for $X$ a discrete random variable with $\lambda > 0$ is defined as:

$$P(X = x) = \frac{e^{-\lambda}\lambda^{x}}{x!} \quad [1]$$

where,

- $e$ is Euler's mathematical constant. $e \approx 2.71828\ldots$
- $x$ is the number of occurrences. $x = 0, 1, 2 \ldots \infty$

As our ML methods assume Gaussian residuals with homoscedastic variance, it is important to pre-transform the data to achieve this.

***B.1. Anscombe transformatiom.*** The Anscombe transform (2) is a statistical data variance stabilization transformation, i.e. a transformation which aims to change the input data so that the variance of each data point or observation is not related to its magnitude. This transformation therefore transforms heteroscedastic Poisson (1) distributed data into approximately Gaussian distributed data with a homoscedastic standard deviation of one. This then allows us to appropriately apply ML algorithms that assume Gaussian variance, which is the vast majority of them. We consider a set of values $x$ in a data set, the Anscombe transform $A$ of x is defined as:

$$A(x) = 2 \times \sqrt{x + \frac{3}{8}} \quad [2]$$

***B.2. Log transformation.*** We want to detect relevant bursts of activity as they may contain key information. In our use case, the percentage change in activity is more important than the absolute change, i.e. an increase in activity from 1 to 2 counts (100% change) is far more important than from 100 to 101 counts (a 1% change). The Logarithm function is a mathematical device to turn multiplicative change into additive change so that the ML can consider multiplicative effects symetrically. Note that applying the Log transform on the output of the Anscombe transform does not cause problems because it is akin to a simple division by two, the Log of $x$ at the power 1/2 is half of the Log of $x$: $\log(x^{1/2}) = \frac{1}{2}\log(x)$.

**C. Need for data imputation.** Visualisation of the raw unfiltered activity data in Fig. S4 reveals the main problems. For both farms for the entirety of the study from January 2015 to April 2016 for the sheep farm and March 2012 to December 2013 for the goat farm, a number of the animals in the herds were equipped with transponders, on the goat farm in February 2013 the transponders were replaced with a newer version, this event is taken into account in order to not mix the data of different animals. The heat maps show the herds where the x axis display the time and the y-axis the transponder unique identification number, each row contains the corresponding transponder activity count data, the pink color highlights missing activity data points. The animals are kraaled (i.e enclosed) every night and released to the pasture every morning, as part of normal husbandry practice and for health evaluation/monitoring the animals are periodically individually inspected after being guided to an inspection facility. From this visualisation we can observe multiple transponders with no or near to no data points likely due to hardware mis-use or failure but also large time frames where the data is missing for the entire herd likely due to the data administrator forgetting to download the data from the servers. In addition, by zooming into the time axis to a day, in Fig S3A we can see missing points likely due to drop of information packet during communication of the transponder and the base station (Fig. S1), packet drop are cause by the transponders being out of range, in Fig S3B we can see that a decrease of transponder signal strength (i.e the amount of energy the radio pushes into the air over time) correlate with a missing activity point, in other words, when the signal strength decrease too much no activity points are transmitted or recorded. Missing points can also be caused by the base station not being able to process large amounts of incoming data simultaneously.

The heat maps in Fig. S4 show the raw data collected over the study period for the Sheep farm Delmas and the Goat farm Cedara. Each heat map contains 1 week of activity data for 13 and 73 transponders for Delmas and Cedara respectively. The heat map grids are configured so that each row of heat maps represents 2 successive weeks of data from the start to the end of the study. The content of each individual heatmap shows the output (activity count) of each sensor aggregated at a 10 minutes bin, i.e., each bin contains the sum of 10 activity counts as the transponders were set up to a minute time interval.

**A** – Activity raw counts

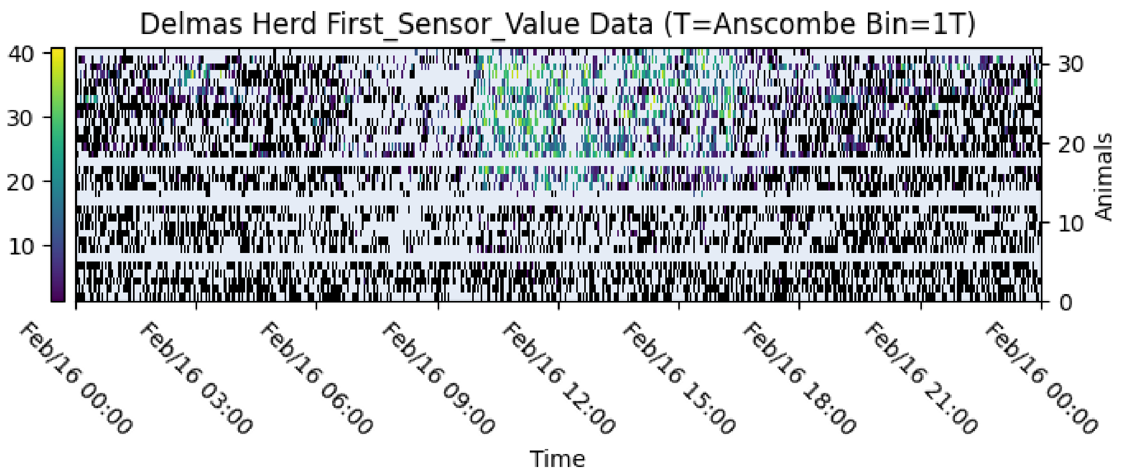


**B** – Signal strength

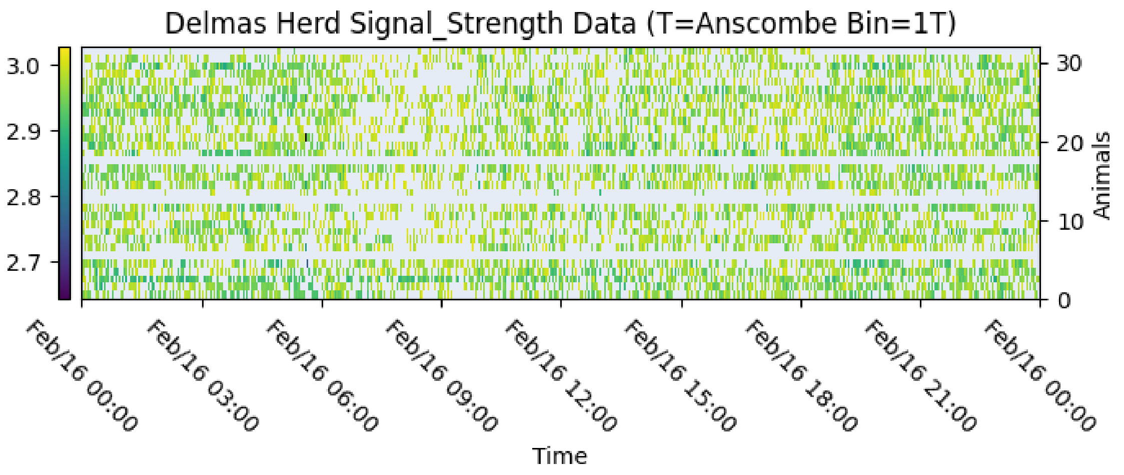


**Fig. S3. Day of raw herd data**. Heat maps of a subsection of the unfiltered raw data, while (A) shows the activity count of multiple transponders (y-axis) after Log and Anscombe transformation, (B) shows the corresponding Logged transformed absolute value of the signal strength of the transponders. Note that the signal strength is measured in dBm (Decibel per milliwatts ranging from 0 to -100), because we are showing the absolute value of the signal strength which is a negative value, higher values indicate to lower signal strength while lower values correspond to high signal strength.

**A Cedara Goat herd activity**

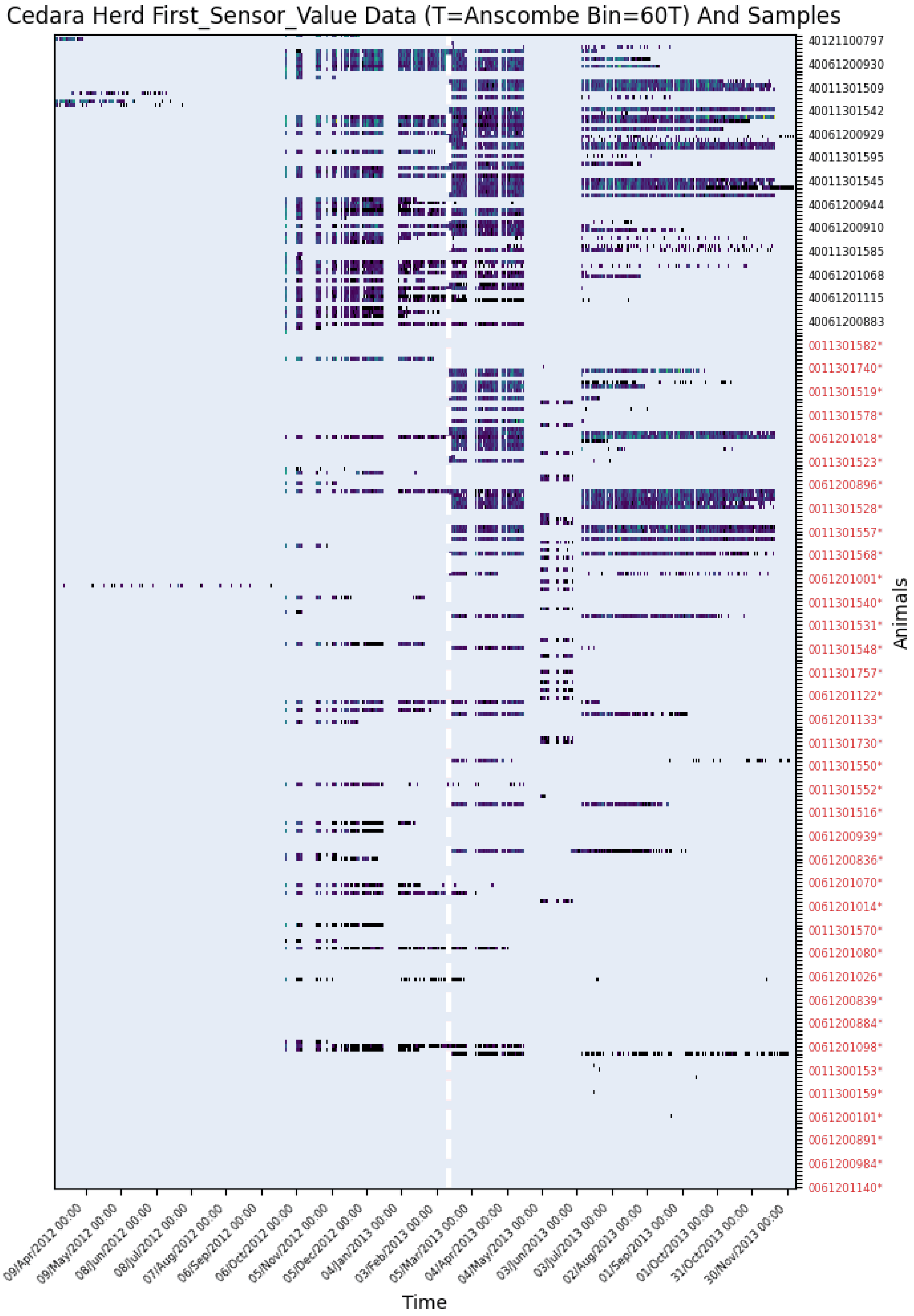


**Fig. S4**

**B Delmas Sheep herd activity**

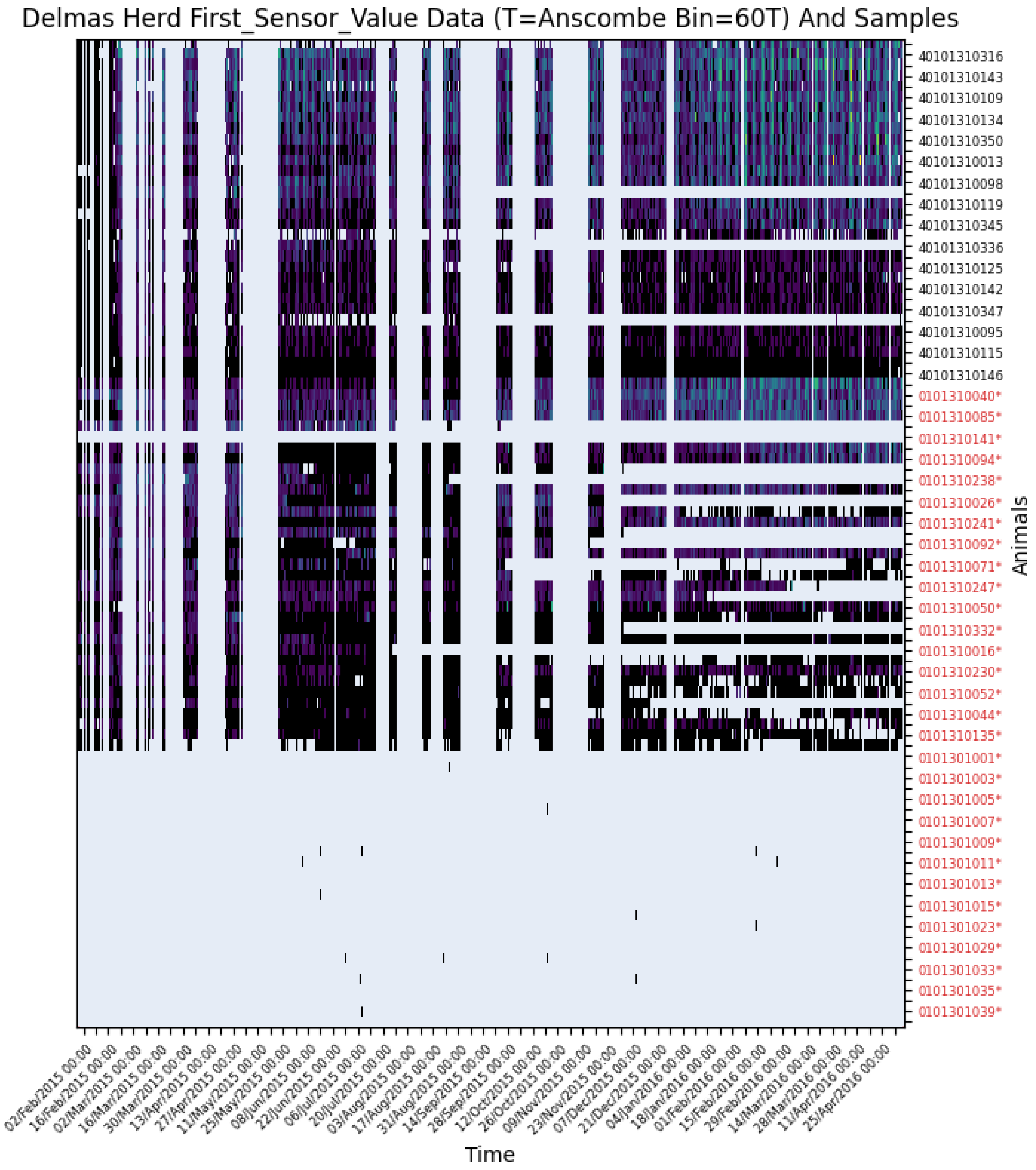


**Fig. S4. Study group herd raw unfiltered activity**. Illustration of the raw activity counts of all of the transponders (y-axis) in the herd, the x-axis shows the time. The pink colour represents missing data points. While (A) shows the transponders data in the goat farm (B) shows the sheep farm. The vertical white dotted line in (A) marks the date of a transponder swap event on the farm. The transponder on the y-axis in red font are transponders with no ground truth data which can't be used in the later chapter for our supervised ML pipeline.

In the next Chapter our machine learning pipeline will rely on modeling a day of activity data to predict animal health. The amount of days that do not contain any missing points in our data-set is limited as we highlighted in this section. It is therefore key to have the maximum number of examples of days for the ML pipeline to be able to work optimally, hence the need for data imputation.

### 2. Data imputation

**A. Introduction.** Missing values are a reality of modern datasets and a common occurrence in data science. They are caused by a variety of reasons ranging from human error to failure of measurement tools, for example in the medical field it is not uncommon for patients to miss appointments or stop their participation to experimental trials, a data set that logged such patient medical record would have missing record. Data imputation is the process in which we infer the missing values. There are many different ways to impute missing data points from very simple approaches such as padding, i.e replacing missing points with a fixed value or more complex techniques that use sophisticated algorithm to estimate missing points. Ultimately we want to replace the missing values with a value that could be true or close to the real value that was missed, for that reason more advance imputation are usually preferred depending on the use case. We will use simple and more advanced methods to impute our data; linear interpolation is a natural simple approach for time series data as it based on a first order approximation, the equation of a line $y = mx + b$ where $m$ is the slope and $b$ is the intercept, both can be calculated if we have at least 2 data points, thanks to the formula we can estimate any missing data points, the points will be said to be "linearly interpolated". This approach is fast but assumes that the missing points depend linearly on the nearest neighbour non-missing points from the same transponder, and are also de-correlated with all the other transponders.

Fang et al (3) surveyed current (2020) time series data imputation methods and found that deep learning approached based on the Recurrent Neural network algorithm (4) outperformed other approaches. For the more complex methods, we will use two different approaches both based on deep learning algorithms that aim to estimate the conditional distribution of the data across the herd.

***A.1. Generative Adversarial Imputation Nets (GAIN).*** GAIN is a missing data imputation technique based on Generative Adversarial Networks (GAN) (5), (6). GANs are composed of two artificial intelligence models which work against each other. Most supervised machine learning models are used to generate a prediction. Classically we start by feeding a model input training data, the model, in turn, outputs a prediction, and we can compare the predicted output with the expected output based on the prediction results we can update the model to improve its predictive power. GAN is an unsupervised technique which consists of two sub-models, a Generator and a Discriminator. The Generator's job is to create fake samples while the Discriminator's job is to take a given sample from the Generator and determine if the sample is fake or real from the original input domain set. As an example to illustrate how GAN works, we might want to generate pictures of human faces. We are aiming to train the Generator to create convincing face pictures. We start by first training the discriminator model to recognize what a picture of a face looks like, hence we need a large data-set of face pictures that we feed into the discriminator which will look at all the attributes that make up a face in order to characterize what a typical face should look like, once the discriminator is capable to recognise real faces then we can test it by feeding it pictures which are not pictures of faces to make sure it can make the difference. During this first phase, the Generator is frozen but once the discriminator gets good at recognizing pictures from the original data-set i.e pictures of faces then we apply the generator to start creating fake pictures of faces, by selecting a random sample from the data-set and modifying it to create a fake sample, the next step will be to feed the discriminator the newly created fake sample and check if it is capable to identify the sample as fake, based on the answer either one or the other sub-models will be updated. If the discriminator successfully spots that the sample is fake then it remains unchanged while the generator will need to update its model to generate better fakes, and vice-versa if the generator creates a fake sample that fools the Discriminator, the discriminator model needs to update itself. We repeat the process until the generator gets so good that the discriminator can no longer pick out its fakes.

GAIN operates on tabular data, using a fully connected neural network (Fig. S6) for both the generator and the discriminator to model the conditional distribution between the columns, while assuming the rows are independent and identically distributed. In addition the generator is given a hint matrix containing the location of the missing data points along side the input data data that we wish to impute, this ensures that the model understand the true distribution of the data, ie the model has a concept of the distribution of the real data and the missing points, (5).

***A.2. Multi-directional Recurrent Neural Networks (M-RNN).*** In our study, we can take advantage of imputation techniques such as the M-RNN approach devised by Yoon et al (8) which takes advantage of the temporal component across multiple data streams. In our use case, this approach can take advantage of the information contained in all of the transponders in the herd to impute the data because it can model both temporal correlations within each transponder and an orthogonal fully-connected neural network to model correlations across transponders.

Recurrent neural networks (4) RNN are a deep learning technique which predict sequences of data, which is commonly used in sequence modelling problems such as natural language processing and stock prediction where the sequence of the data is important. Traditional RNNs are also called feed-forward neural networks and use the concept of sequential memory by using previous information to affect later ones, this takes the form of a looping mechanism that allows information to flow from one step to the next, and previous information is saved in a hidden state.

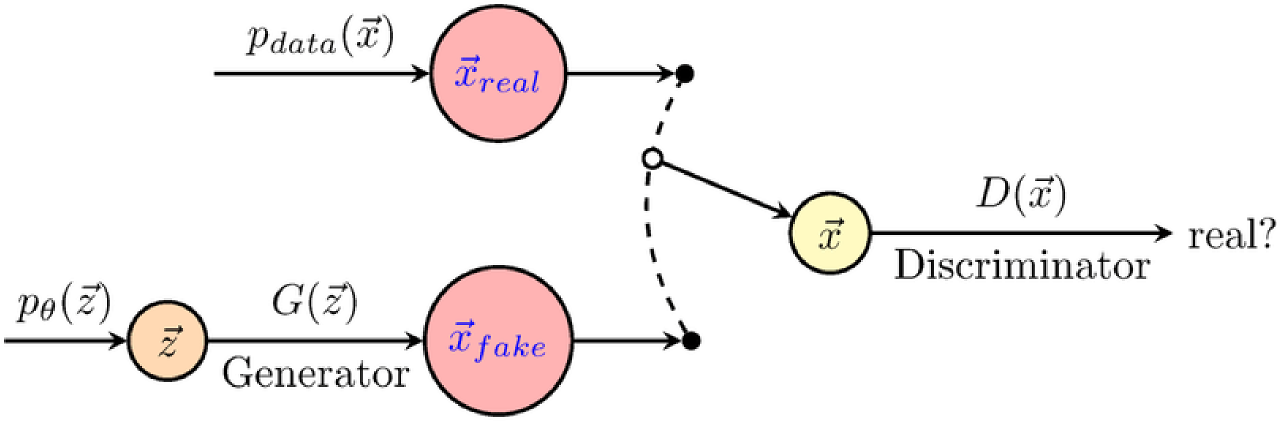


**Fig. S5.** Schematic of generative adversarial networks (GANs). The generator (G), defines a probability distribution based on the information from the samples, whereas, the discriminator (D) distinguishes data produced by G from the real data. Figure original source: (7).

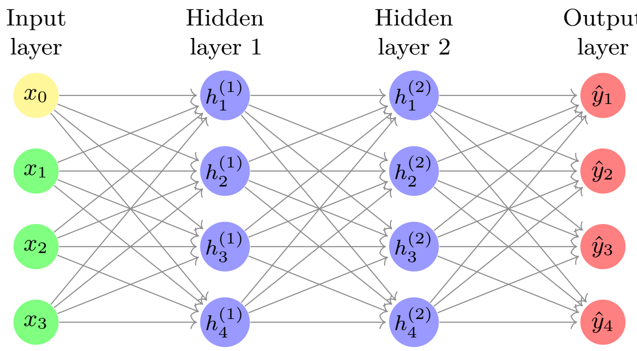


**Fig. S6. Schematic of a Fully Connected Neural Network**. (Generated with the tikz library, author:Till Tantau)

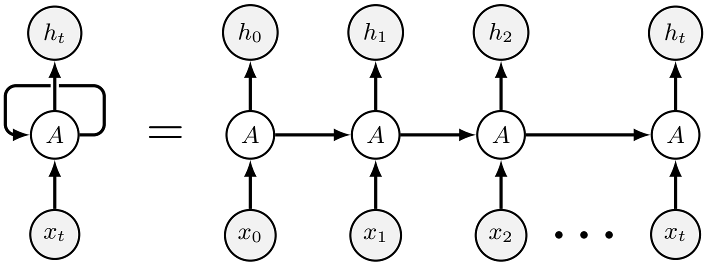


**Fig. S7. Recurrent neural network diagram** $x$, $A$ and $h$ are the input, hidden and output layers respectively during the different learning steps, ie for each element in the input sequence $x$. (Generated with the tikz library, author:Till Tantau)

Short term memory is a known limitation of RNNs, it is caused by the vanishing gradient problem (9) which is due to the nature of back-propagation (10), training a Deep Neural Network has 3 major steps, first, a forward pass that makes a prediction, second, a comparison of the prediction to ground truth using a loss function which output an error value which estimates how "badly" the network is performing, last the error value is used to calculate the gradient in each node in the network through back propagation, the gradient being the value which is used to calculate the network internal weights which ultimately allows the network to learn, the scale of the adjustment to the internal weights of the network is directly linked to the gradient value, the larger the gradient the larger the weight adjustments will be, similarly small gradient values will cause small adjustments. When doing back-propagation each node calculates its gradients with respect to the gradients from the nodes in the previous layer which is a problem as the gradient in the next layer would be smaller after each consecutive layer preventing the earlier layer from "learning" has their weight are only adjusted by a very small amount due to small gradient values.

In an RNN each time step is a layer, to train an RNN back-propagation through time (11) is used. The gradient values will exponentially shrink as it propagates for each time step, which renders the model unable to learn from earlier time steps which are vital in most sequence modelling problems, for example, to predict the next word in a sentence knowing a large number of words is crucial for context, basing the prediction on just a few latter words can cause out of context predictions. To combat the short term memory problem of RNN, two main specialized RNN was created, Long Short Term Memory (LSTM) (12) and Gated Recurrent Units (GRU) (13) which are capable of learning long-term dependencies using mechanism called gates which are different tensor operations that can learn what information to add or remove to the hidden state.

MRNN is composed of multiple independent bidirectional RNNs (Fig. S7), each model the data in an individual stream of data, in our case individual transponder data. In addition, a Neural Network fully connected (FC) layer (Fig. S6) is used to model the data across every transponder, where each layer in the FC layer contains the data of a single transponder. This last model can model important data from the herd which improves imputation results at the animal level (C).

### B. Methods.

***B.1. Selecting high-quality training data.*** It is important to train our imputation models with high quality data that reflects best the real world activity of the animals. For this research the data collection not only suffered from missing data points but also invalid and weak signals due to the transponders not being used correctly or placed on the animals securely, for example some workers have reported to leave active transponders on a desk for a prolonged amount of time, such events were not logged, for that reason we will analyse the quality of the data to assure that the data that we used for this research is valid. For each transponder, we evaluate the complexity (14) of the accelerometry data with the Shannon entropy (15), in order to select the transponder that contains the densest information and dismiss the lower quality (i.e, large amount of missingness and abnormal data points) traces. The Shannon Entropy can help us measure the information content in the transponder which ultimately allow us to detect invalid transponders. The two main problems the transponder traces suffer from are repetition of the same activity count value for an extensive period of time and excessive amount of missing points. This approach is effective at detecting both. The Shannon Entropy formula is given by:

$$H = -\sum_{i=1}^{M} P_i \, log_2 \, P_i. \qquad [3]$$

Where $P_i$ is the normalised probability distribution of the count in a transponder time series.

**A** – Sheep farm transponders sorted by increasing entropy

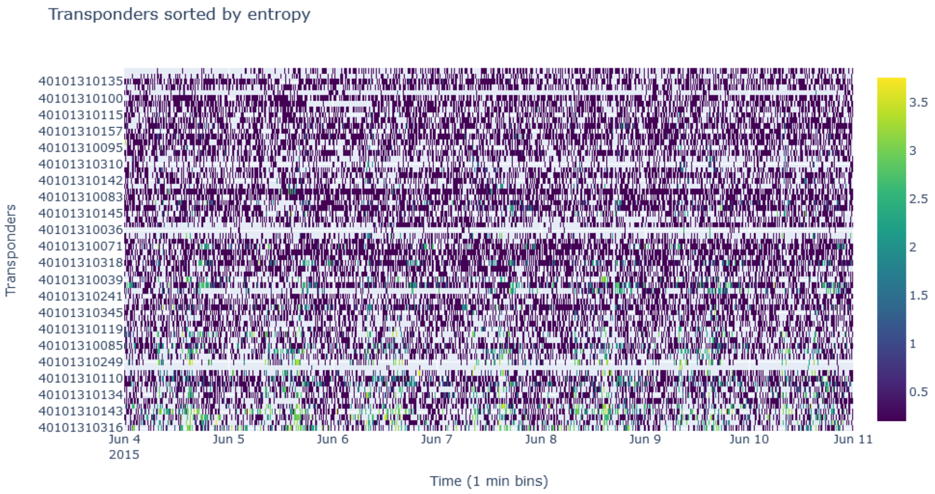


**B** –Goat farm transponders sorted by increasing entropy

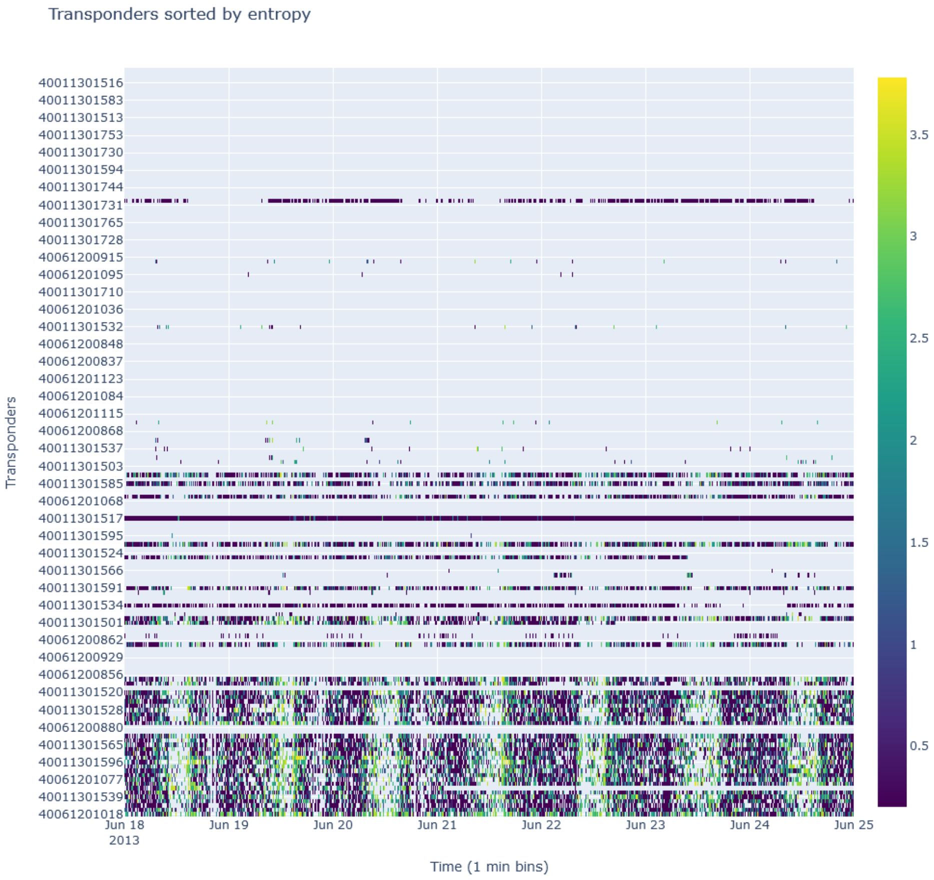


**Fig. S8. Illustration of transponder data sorted by entropy**. This heatmap shows a 7 day section of all of the transponders sorted with their respective Shannon entropy in ascending order from the bottom to the top of the y axis. The raw activity counts are Anscombe and then Log transformed in this visualisation. We can observe the improvement of the "quality" of the transponders with lower entries. It is important to note that this illustration only shows a subsection of the time frame, transponders with no data in this subsection do not necessarily have a lot of missing points in other time frames. Missing points are displayed via the light turquoise color.

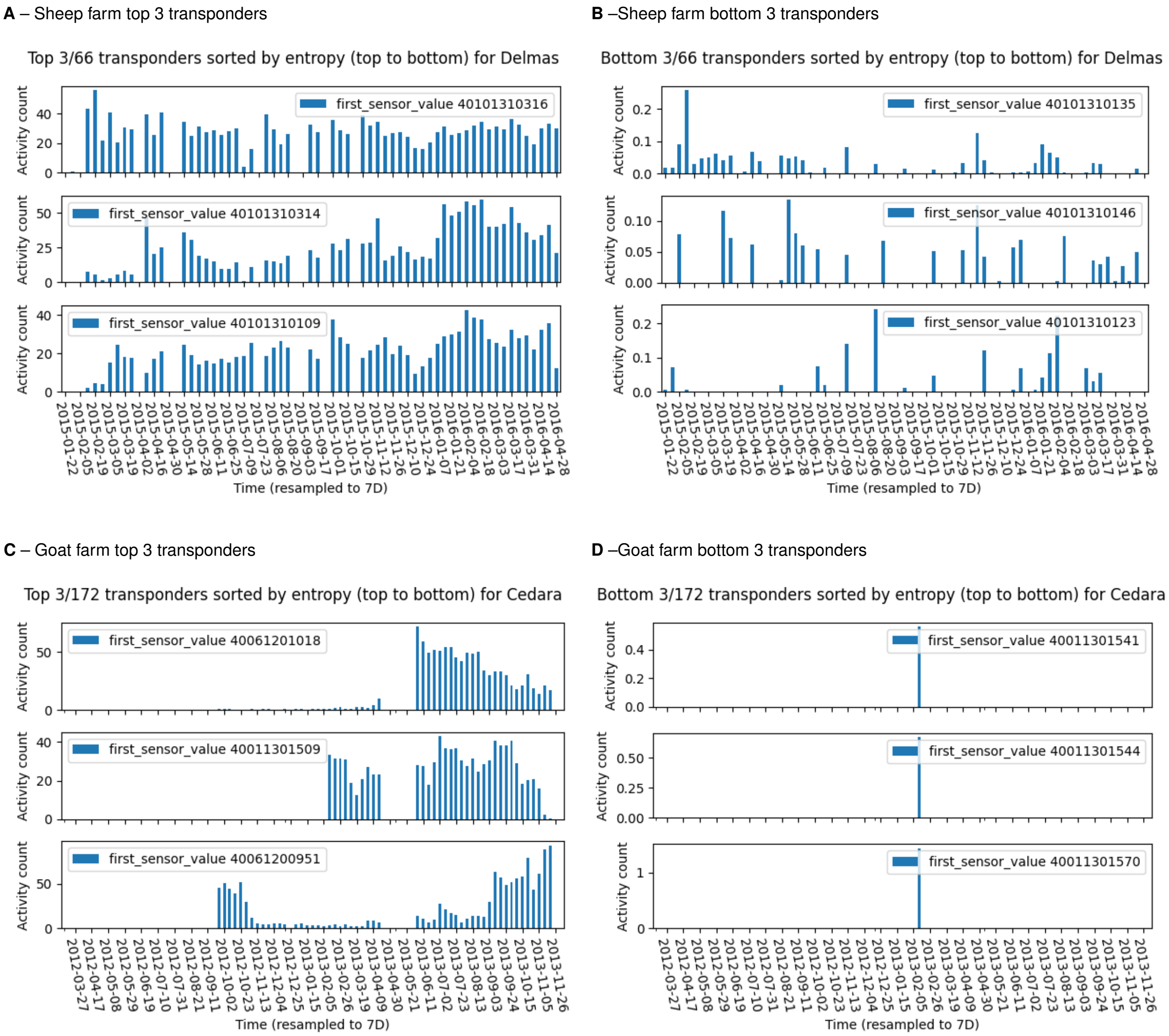


**Fig. S9. Overall view of transponder data quality**. This figure shows the top 3 and bottom 3 transponders based on their respective data entropy. For visualisation convenience we re-sampled the data to a 7 days resolution by summing all data points within a week. While (A) and (B) show the Sheep transponders (C) and (D) show the Goat transponders.

***B.2. Creating input samples.*** Here we will illustrate with a simple example how we will create the input samples for the GAIN and M-RNN techniques (Fig. S10). In this example the accelerometry data come from 2 different transponders, each collected twelve data points over time, transponder 1 could not measure accelerometry data 4 times at time $t4$, $t9$, $t11$ and $t12$ while transponder 2 only missed the data point at time $t5$. In both case we need to introduce the concept of window, i.e. a subset of the original input data of a variable length, in our example we used a window length of 4 points (in our study the data is sampled to a minute resolution, hence 4 points correspond to 4 minutes). For GAIN for each transponder we build samples by sliding the window along its time axis with a stride equal to the window length, in that fashion we can build 3 samples per transponders for a total of 6 samples for the GAIN algorithm. In comparison for M-RNN we slide the widow along the time axis common to every transponders so that each samples contains the data of all transponders at all times, in our case we could build 3 samples. The key difference is that M-RNN accepts and models a matrix of data for each sample, wherease GAIN only models a vector.

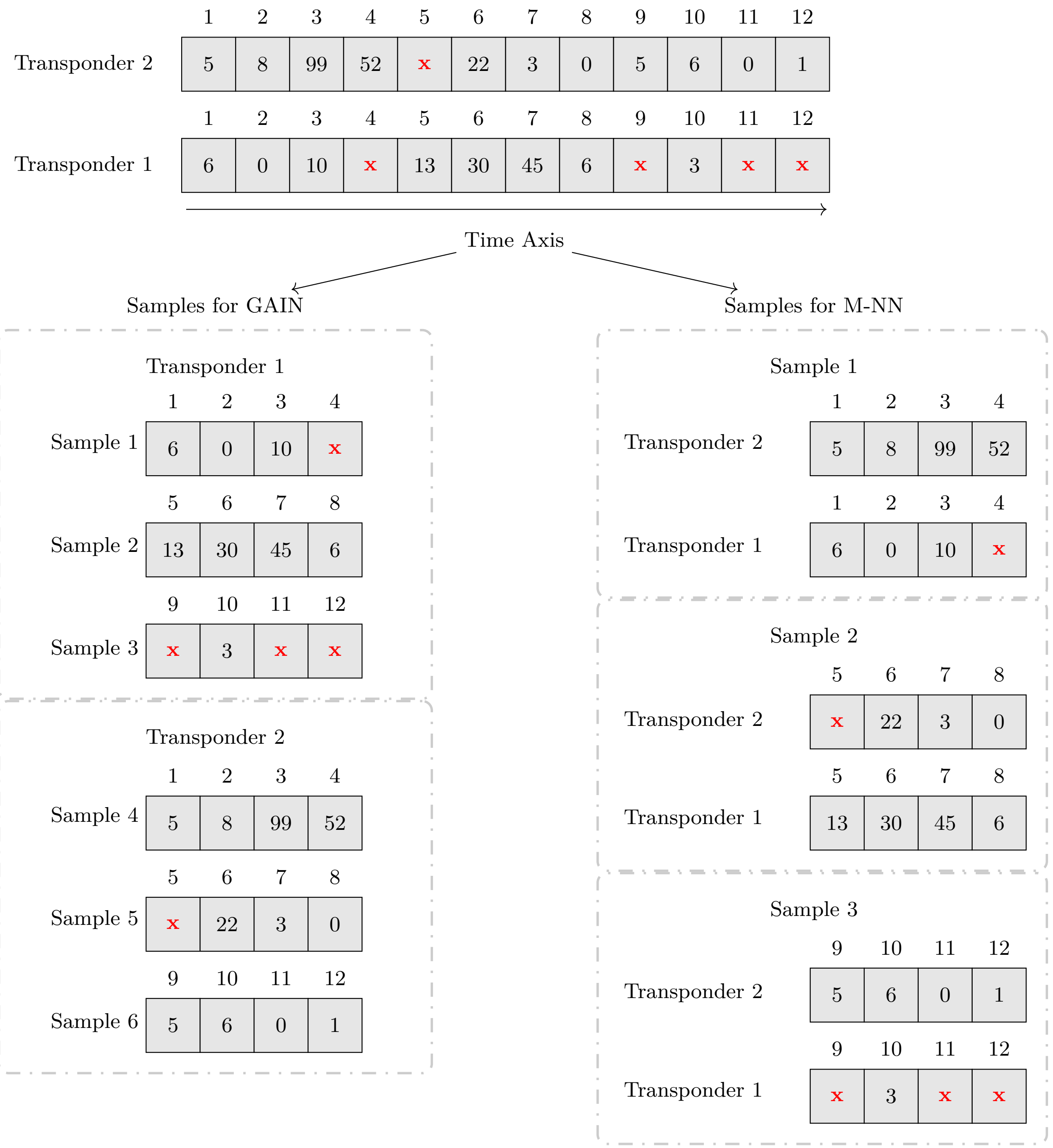


**Fig. S10. Explanation of the data preparation.**This schematics shows how the samples are created for the two main imputation techniques we used. Generative Adversarial Nets Imputation (GAIN) (5) does not not inherently have a concept of the entire timeline of the data stream while Multi-Directional Recurrent Neural Network (M-RNN) (8) natively uses the timeline, the natural temporal order of the samples is taken into consideration.

The pipeline to implement the GAIN and M-RNN algorithms are illustrated in Fig. S11 and explained in detail in the following subsections.

***B.3. GAIN implementation.*** For our application, the aim of GAIN is to create convincing fakes of our activity samples which can be used to impute our data set. We used the implementation developed by Yoon et al (5) and adapted to our use case. Below, we describe the different steps (Schematic. S11) to impute the missing points in the original accelerometry traces. We aim to train the GAIN model with the accelerometry data recorded during a single or multiple days, which will allow us to capture and model the activity data of the selected time frames, we will investigate the optimal length or time frame to use given our data set.

We consider a data set of $N$ animals. For each animal $a$ we have a data stream of length $T$ where each data point is measured at the time $t$.

$$Input(N,T) = \begin{bmatrix} a_0(t_0), \dots, a_0(t_T) \\ \ddots \\ a_N(t_0), \dots, a_N(t_T) \end{bmatrix} \quad [4]$$

We want to train a GAIN with samples that contain accelerometry data and other features $F$ that can help the algorithm to optimally model the data, in our case we want to model what a day of activity looks like. $F$ is an array that holds the timestamp of the day and the unique transponder id associated with the accelerometry data. By stacking days of activity data from multiple transponders GAIN can learn across the entire herd what a typical day of activity looks like.

$$samples(N,T) = \begin{bmatrix} a_0(t_0), \dots, a_0(t_{1440}), F_0 \\ a_0(t_{1440}), \dots, a_0(t_{2880}), F_0 \\ \vdots \\ a_0(t_{T-1440}), \dots, a_0(t_T), F_0 \\ \vdots \\ a_N(t_{T-1440}), \dots, a_N(t_T), F_N \end{bmatrix} \quad [5]$$

- Training
  1. Raw data cleaning (Section. A).
  2. Read each transponder file, and filter out the transponder with less than 100 non-missing points.
  3. We sort transponders by entropy.
  4. We calculate the log (B.2) of the Anscombe (B.1) of every data point.
  5. We chunk all transponder traces into n-day-length chunks and append the day timestamp and the transponder id for each chunk. (Eq. 5)
  6. We filter out days where more than 80% of the data points are missing. There is 1 accelerometry data point per minute, therefore 1440 points in a day, if more than 1152 points are missing the chunk is dismissed.
  7. The data is normalised in the range of 0 to 1.
  8. The empty data points are masked and replaced with 0.
  9. We can finally train the GAIN model with high-quality n-day-length samples (Fig. S20).
- Testing
  1. We reshape the accelerometry traces we wish to impute into n-day-length chunks.
  2. We apply the same preprocessing approach used during training, namely calculating the Log(B.2) of the Anscombe (B.1) of all data points.
  3. We can feed the trained GAIN model with the n-day-length accelerometry trace to impute the missing point.
  4. We now back-transform the data point into accelerometry counts by calculating the inverse of the Anscombe and Log transform and reverse Normalisation from the range 0 to 1.
  5. finally we restore the imputed data into its original shape of 1 accelerometry trace per transponder.

***B.4. M-RNN implementation.*** In practice, M-RNN takes a list of 2D time series as input, in our case each time series correspond to the raw count activity data of the transponders in the herd. We consider a dataset of $N$ animals. For each animal $a$ we have a data stream of length $T$ where each data point is measured at the time $t$ (Fig. S14). Compared to GAIN where each transponder is treated as an independent data stream, M-RNN uses groups of sensors aligned on the same time axis, in other words, the herd activity needs to be preserved to benefit from this technique. Based on the amount of available data, we aim to model what a week of herd accelerometry data looks like, the fitted model would then be able to impute and predict what a week of accelerometry data should be based on the entire information contained in the herd data.

$$HerdData = \begin{bmatrix} a_0(t_0), \dots, a_0(t_T) \\ \ddots \\ a_N(t_0), \dots, a_N(t_T) \end{bmatrix} \quad [6]$$

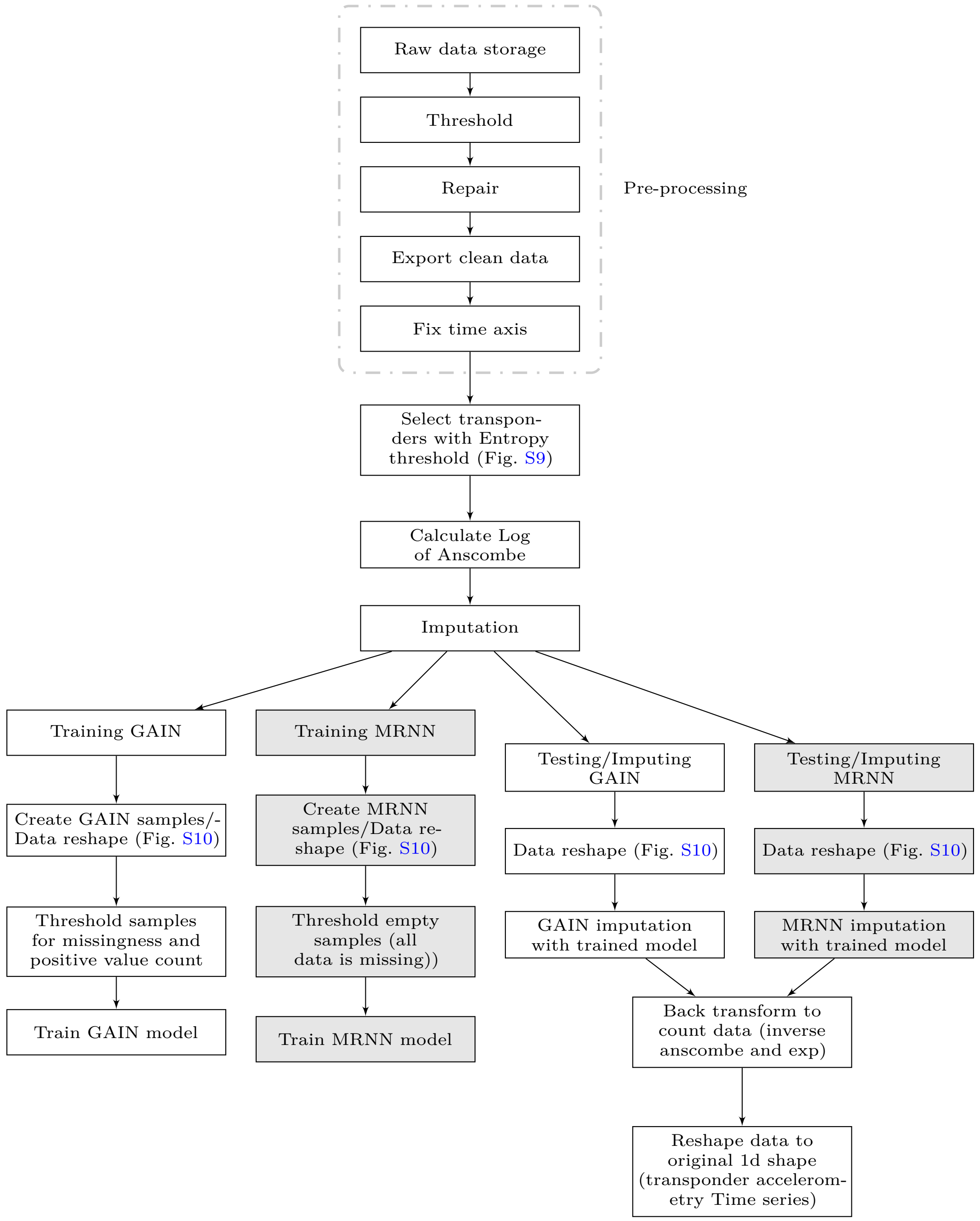


**Fig. S11. Schematic of the GAIN and MRNN imputation pipeline**. This flowchart illustrates the key steps for each imputation approach.

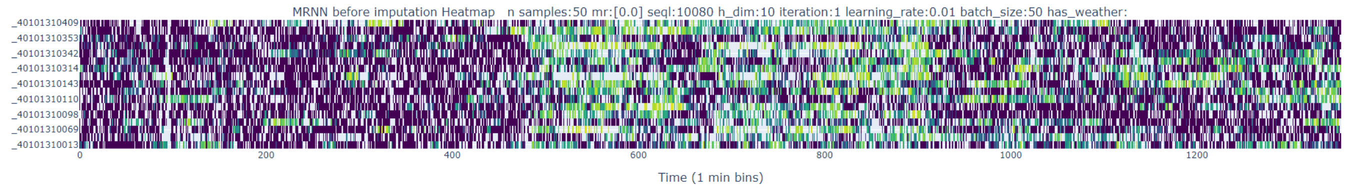


**Fig. S12. This figure shows a subset of the accelerometry input data for M-RNN** each row shows a 1440-minute long stream of activity data after Anscombe and Log transformation where each data point is measured every minute.

Because M-RNN uses the mean square error as a cost function we first apply the Anscombe ( B.1) and the Log ( B.2) transformation to make the native Poisson distributed count follows a normal distribution.
To train the M-RNN model training sets are built by creating consecutive subsections of the input data, each $i$ subsection $w(N,t)_i$ is of fixed length $s$ where $t_s - t_n = s$ and $n$ is incremented by $s$. Here $w$ is the accelerometry data component of the sample where each row value comes from a different transponder, each sample is constructed by creating 3D matrix where the first matrix is $w_i$, a second mask matrix that will hold the location of the missing points in $w_i$, and a final matrix that for each row of $w_i$

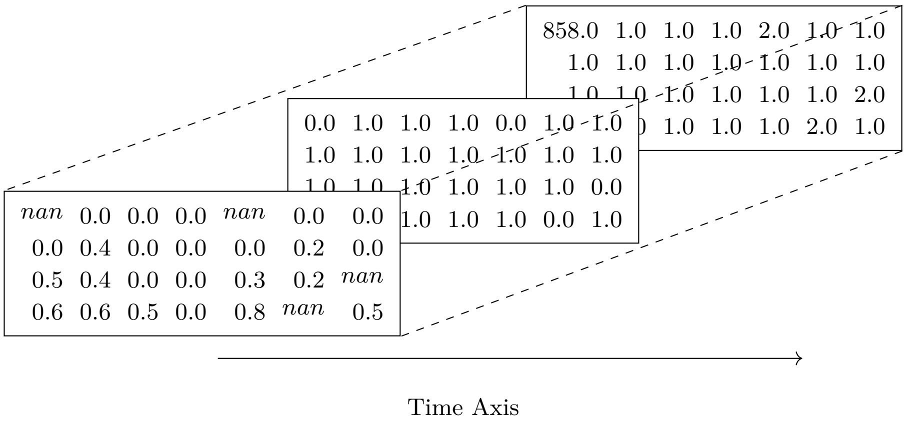


**Fig. S13. Schematic of the M-RNN imputation 3D sample)**. In this illustration, for all matrixs each row corresponds to the data stream from a single transponder, in this example, there are 4 transponders data streams. The columns represent time, there are 7 consecutive accelerometry records here. The first matrix (in the front) shows the accelerometry data, and the missing points are marked by the word 'nan'(i.e not a number). The other matrixs are calculated based on the first. The matrix in the middle is a mask of the first one while the third one shows how many data points were missing before a new data point is recorded by the transponder, in this example the top transponder could not record 858 values before it could record a real data point (0).

$$w(N,t)_i = \begin{bmatrix} a_0(t_0), \ldots, a_0(t_s) \\ \ddots \\ a_N(t_0), \ldots, a_N(t_s) \end{bmatrix} \cdots \begin{bmatrix} a_0(t_s), \ldots, a_0(t_{s*2}) \\ \ddots \\ a_N(t_s), \ldots, a_N(t_{s*2}) \end{bmatrix} \cdots \begin{bmatrix} a_0(t_{s*i-1}), \ldots, a_0(t_{s*i}) \\ \ddots \\ a_N(t_{s*i-1}), \ldots, a_N(t_{s*i}) \end{bmatrix} \quad [7]$$

- Training
  1. Raw data cleaning (Section. A).
  2. We calculate the Log (B.2) of the Anscombe (B.1) of every data point.
  3. The data is normalised in the range of 0 to 1.
  4. We build and reshape the entire data into multiple 3D matrices (activity, mask, time) (Fig. S13) of 7 days long with a sliding window along the time axis, we used a 7 days stride with no overlap.
  5. We filter out the windows that do not contain any activity data.
  6. Fitting all the Bi-RNNs (one per transponder)
  7. Fitting the Fully connected layer
- Testing
  1. Reshape data into 3D matrices (activity, mask, time) (Fig. S13).
  2. We can feed MRNN with the reshaped data.
  3. If data points are missing in all streams impute with the data from the mean window.

4. Revert MinMax Scale normalisation and inverse Log (Apply natural exp) and Anscombe transform
5. Restore original data traces shape

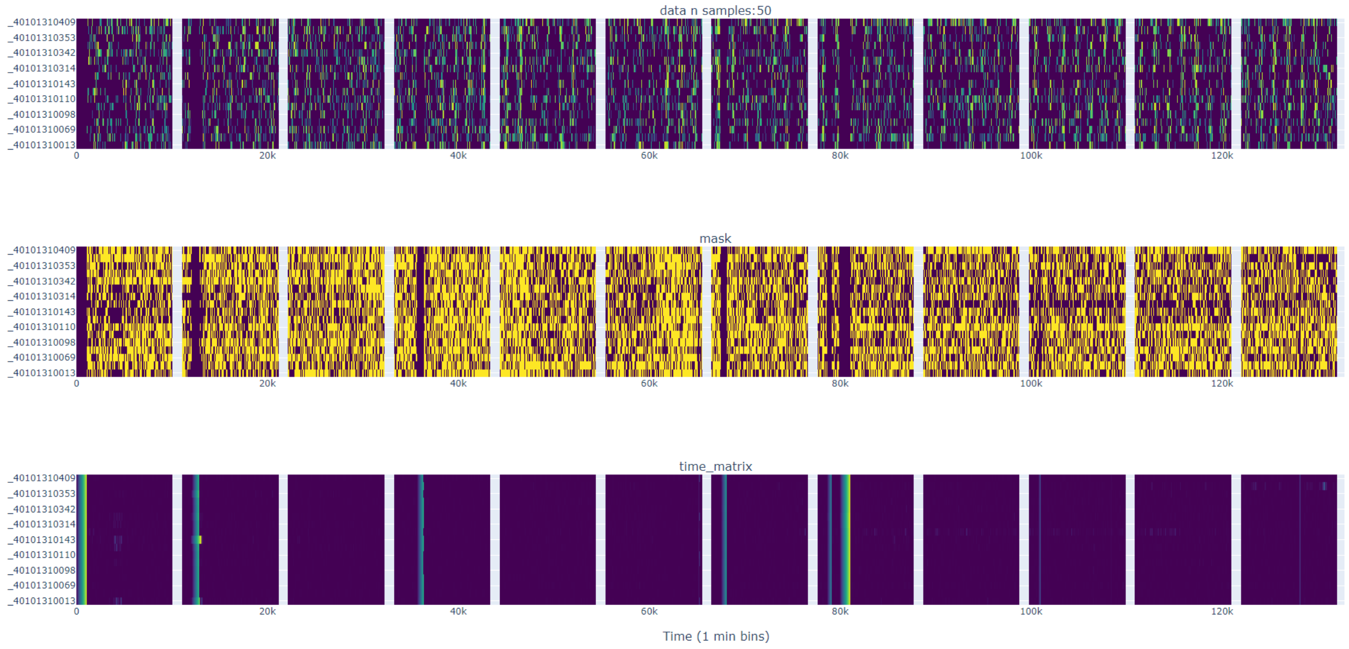


**Fig. S14. Example of the training set for M-RNN** with real Sheep accelerometry data. In this example, we show 12 samples ready for training a M-RNN imputation model. Each column corresponds to the 3 elements of a 3D matrix (Fig. S13) from the top to the bottom, first the activity data, then the mask data, and finally the "time since missing point" data.

***B.5. Evaluation.*** We can evaluate the result from the GAIN and MRNN imputation by adding missing data points and comparing the imputed point with the original value of the same transponders. We will use the Root mean square error (RMSE) (16) for that purpose and we will compare the imputation results with imputed values achieved via linear interpolation of the data on the time axis. Lower RMSE values are better as the indicate that the imputed value is closer to the real value. The experiment shows that the MRNN-imputed data points are closer to their original value compared to the GAIN and linearly imputed one, which suggests that MRNN is more accurate at predicting missing values (Fig. S16). In this section we will present the results of different imputation experiment to establish the optimal methodology and parameters. For the GAIN imputation we will investigate the effects of the filtering threshold, the sample length (window size) and the effect of adding extra features in the training samples. For the MRNN imputation we will experiment with the same parameters but also monitor the impact of using different number of data stream for training.

### C. Results.

***C.1. Robustness toward missingness.*** By adding missing points to the original samples we can evaluate how resistant to missingness the imputation the imputation is. This can also give us a clear indication of which technique is most powerful for our use case. Figure S15 shows that M-RNN performs better for all sample length compared to GAIN.

***C.2. Sample length and RMSE.*** Ultimately we want to find the optimal training configuration for each of the algorithms before comparing their outputs, we can use the RMSE as a metric to determine the best sample length to use. Here will add 40% of missingness, i.e 40% of the dataset points will be marked as missing (Fig S16C).

For the GAIN imputation sample length of 1 day gives us a RMSE slightly below 0.2 while other sample length give values $> 0.2$. Similarly 1 day give the lowest RMSE at $< 0.12$ for MRNN. For both techniques increasing the sample length minimally increase the RMSE. Linear interpolation perform the worst with a RMSE around 0.60 for the lowest level of missingness.

***C.3. MRNN training and RMSE.*** We can train the MRNN model with a subset of high entropy transponders to ensure we are using the best data for training our models, however we need to be cautious about over curating our data as some level of noise is realistic in a on the field scenario, for that reason we will train the models with different number of transponders and evaluate the effect on the RMSE. An added benefit of training on a subset of the transponders significantly decrease processing time as MRNN involves training as many Bi-RNN as transponders used for training.

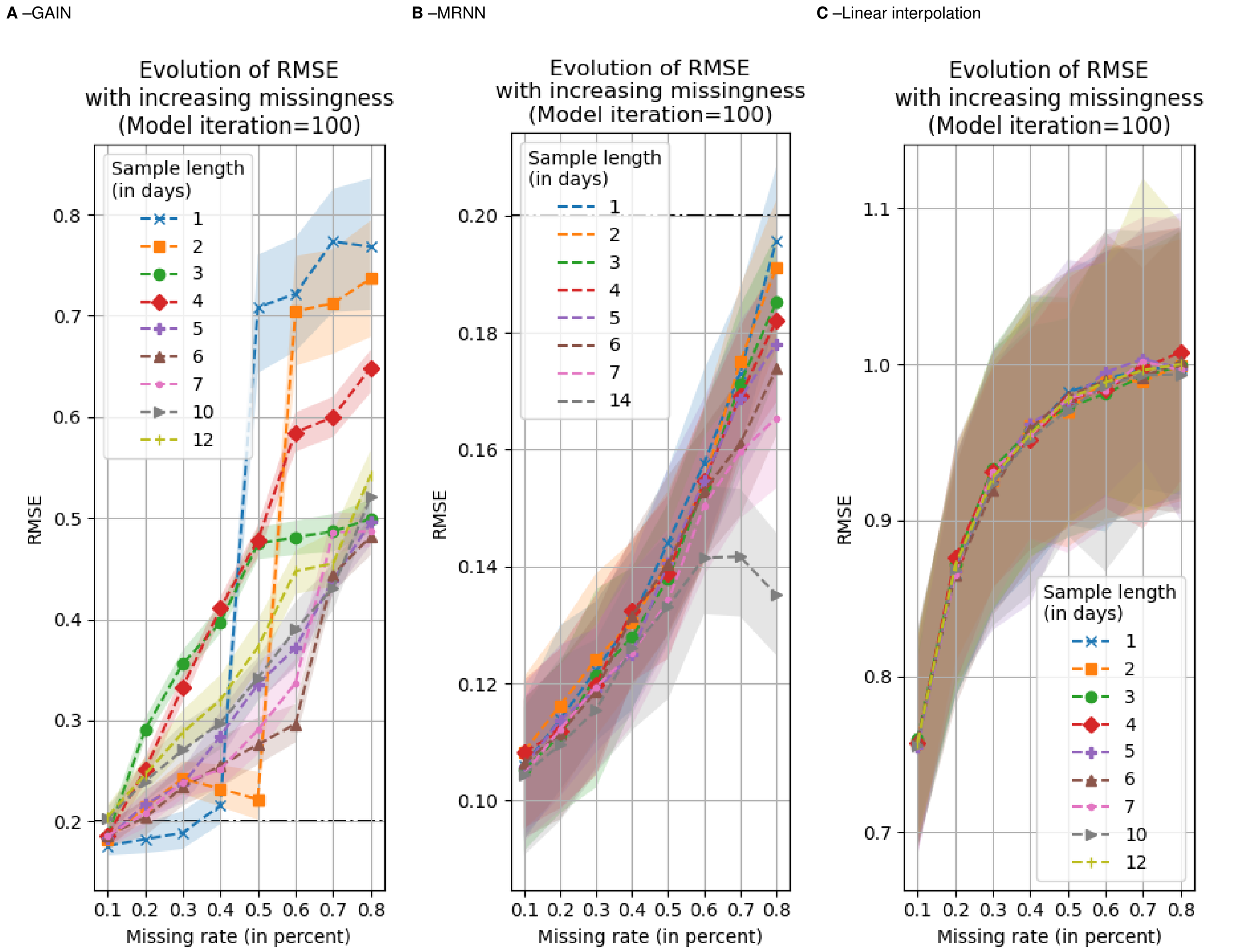


**Fig. S15. Evolution of RMSE with increased missingness for GAIN and MRNN**. The x-axis shows the missing rate, i.e the rate of artificially added missing points. The y-axis shows the root mean square error (RMSE). While (A) shows the results for GAIN (B) shows the results for MRNN and (C) Linear interpolation.

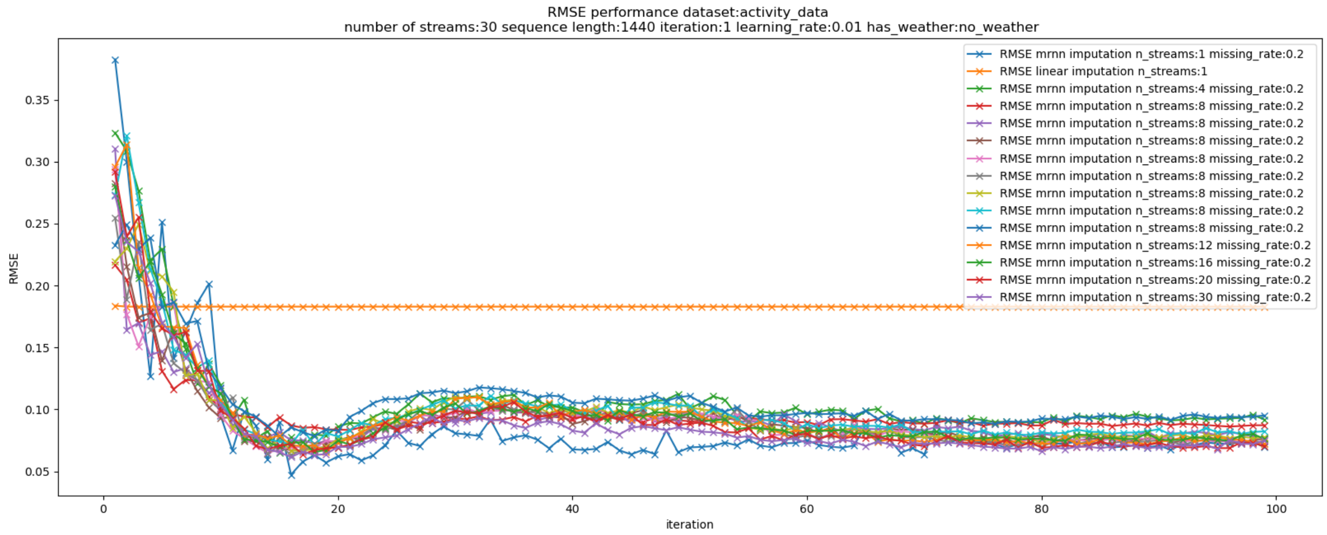


**Fig. S17. Comparison of The Root mean square error (RMSE) of MRNN** with training on different number of transponders (data stream).

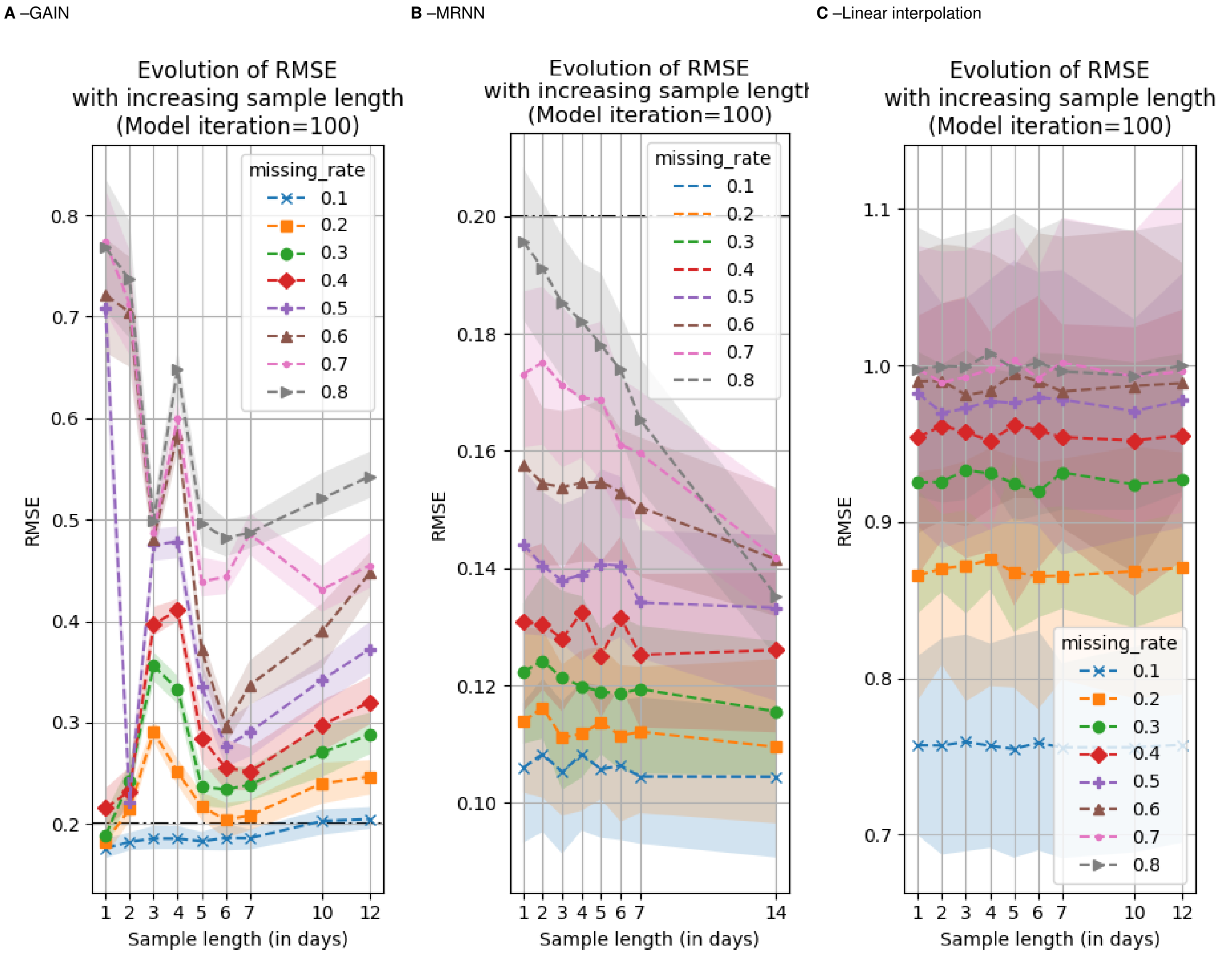


**Fig. S16. Evolution of RMSE with increased sample length for GAIN and MRNN**. The x-axis shows the sample length in days, i.e the number of activity data points in the sample, for example 1-day samples contain 1440 data points, 1 per minute. The y-axis shows the root mean square error (RMSE). While (A) shows the results for GAIN (B) shows the results for MRNN and (C) Linear interpolation.

***C.4. Model loss curves and sample length.*** Analysing the model loss function can give us an indication of how well the model is learning. For the GAIN models we expect the two competing model to diverge as the Generator gets better at crating fakes and the discriminator better at detecting them (Section. A.1). For MRNN we expect the loss of the model to decrease as it is learning.

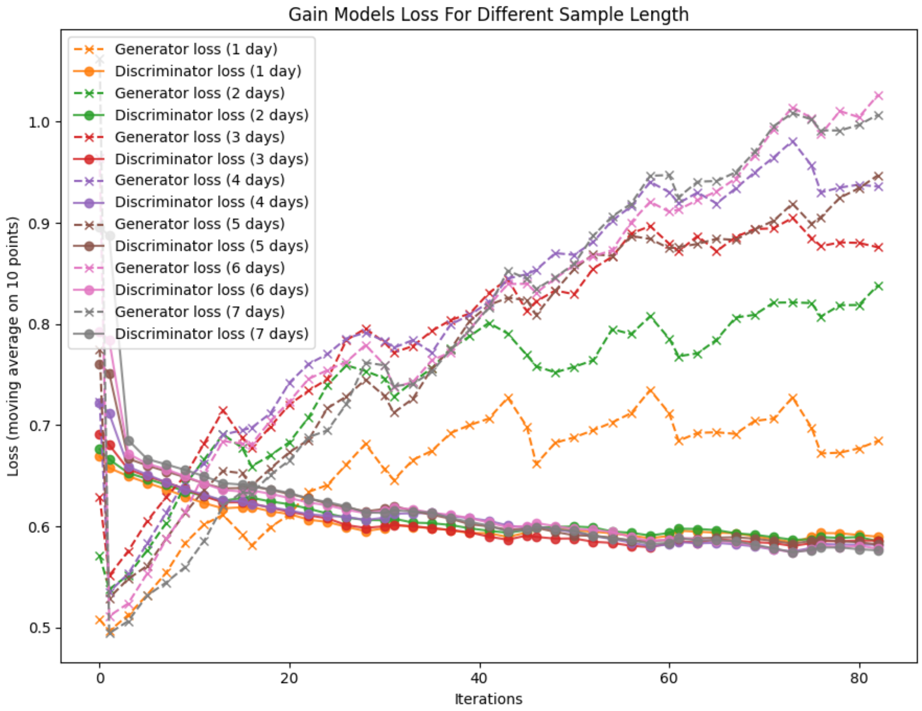


**Fig. S18. Evolution of RMSE with changing sample length.** The x-axis shows the model iteration while the y-axis shows the moving average of the model loss. Each GAIN model (Generator and Discriminator) share the same color, the generator curves are marked with the "x" marker while the discriminators are marked with "●".

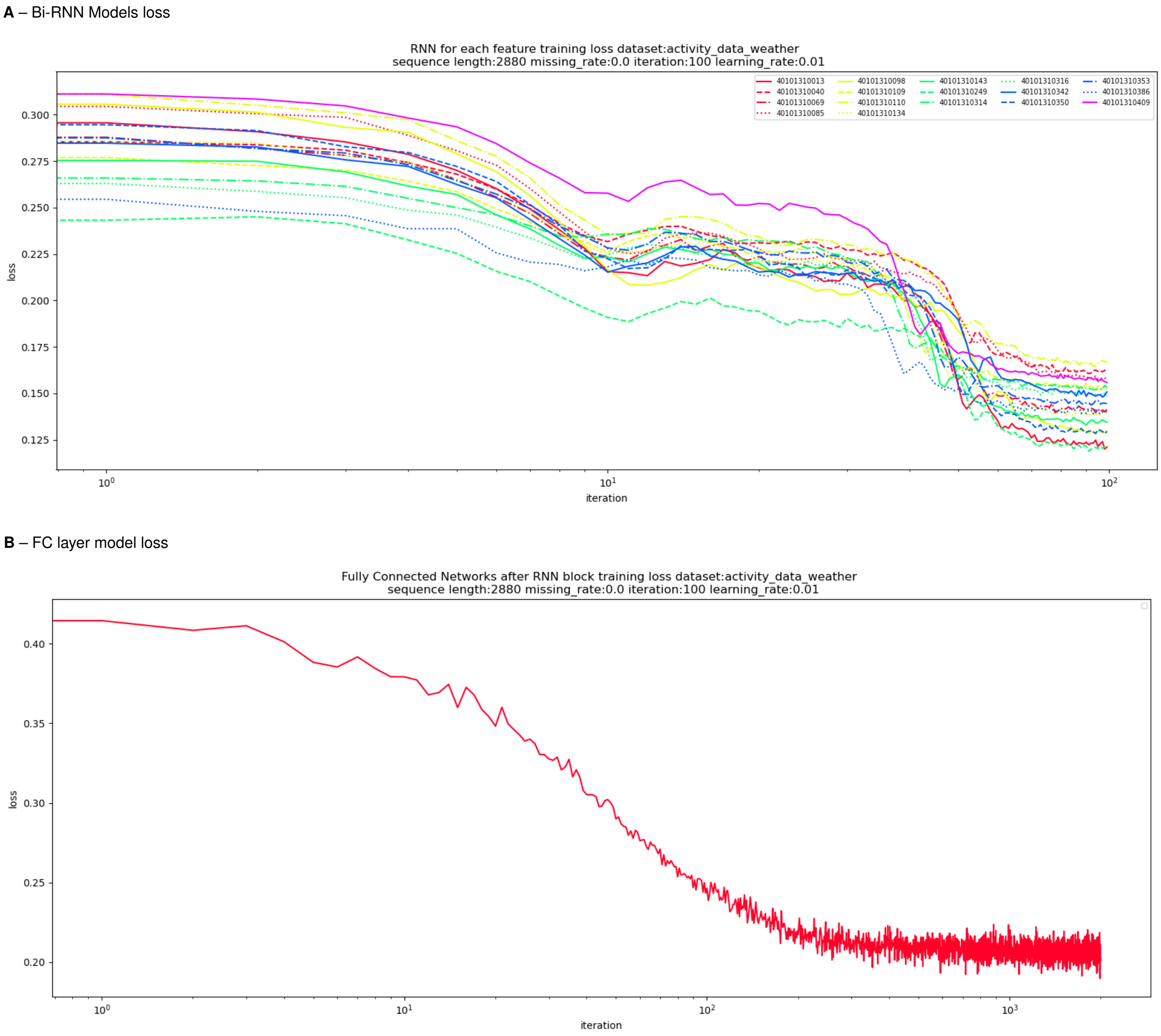


**Fig. S19. MRNN models loss**. (A) shows the loss of each Bi-RNN while (B) shows the loss of the fully connected layer. The x-axis shows the model iteration and the y-axis shows the loss. In (A) each curve correspond to a Bi-RNN model learning the data conditional distribution of a single transponder, we can observe that some model converge faster than others.

***C.5. Imputation visualisation.*** In this section we will display and compare the output of the imputation with the help of heat maps. We will compare the output of GAIN and MRNN with Linear interpolation. In Figure S20C we can observe that GAIN have successfully learned the structure of a day of activity, particularly for the samples range $[0, \ldots, 20]$ which real values are low, in contrast the linear interpolation approach Fig S20D has no concept of a typical day structure. Similarly Figure S21 highlight how well MRNN learned the data structure of each transponder as it can impute entire chunk of missing herd data, for example after 9.5k minutes of the x-axis.

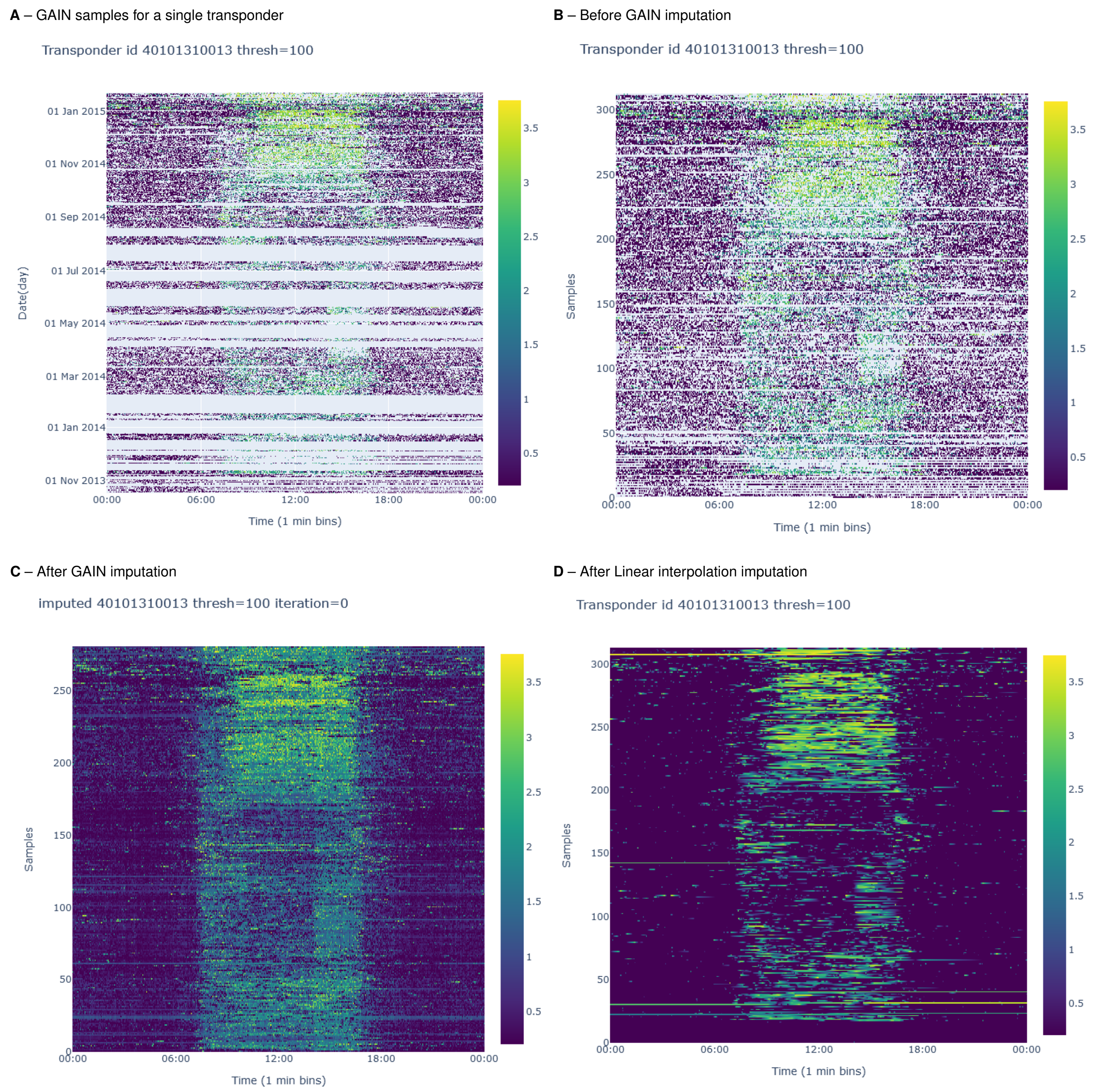


**Fig. S20. Illustration of GAIN training samples from a single transponder for 1 day length samples.** For visualisation purposes, the samples are placed on the entire transponder data collection time frame which displays the period of emptiness in light turquoise (A). (B) shows ready-for-training samples for a single transponder while (C) displays the GAIN imputed output and (D) shows the linear interpolated samples.

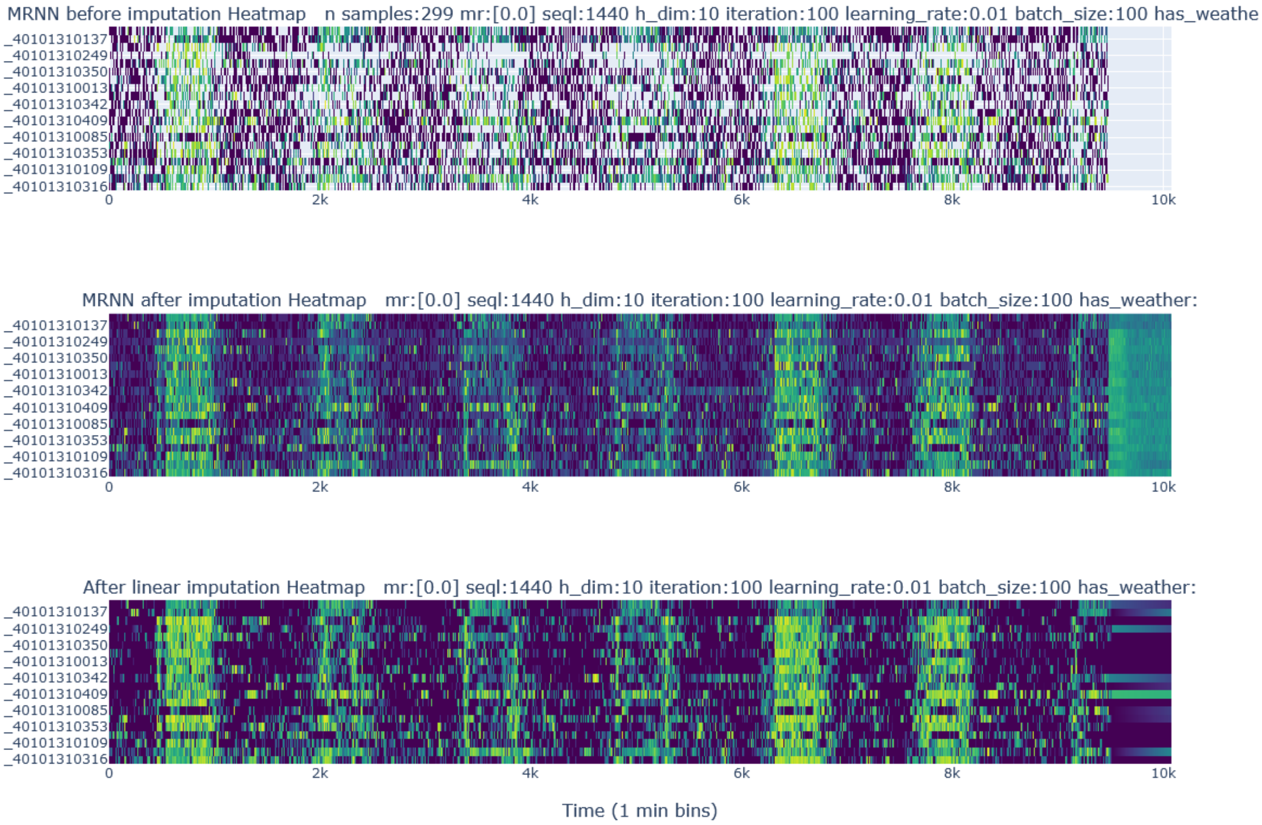


**Fig. S21. Illustration of MRNN imputation on a 7 days subset of herd data.** The top heatmap shows the herd data where each row is the data stream from a single transponder where the missing points are highlighted in light turquoise. The middle heatmap shows the MRNN imputation results while the last heatmap shows the original data imputed with linear interpolation within each transponder trace.

### 3. Building samples for machine learning

**A** – Sheep

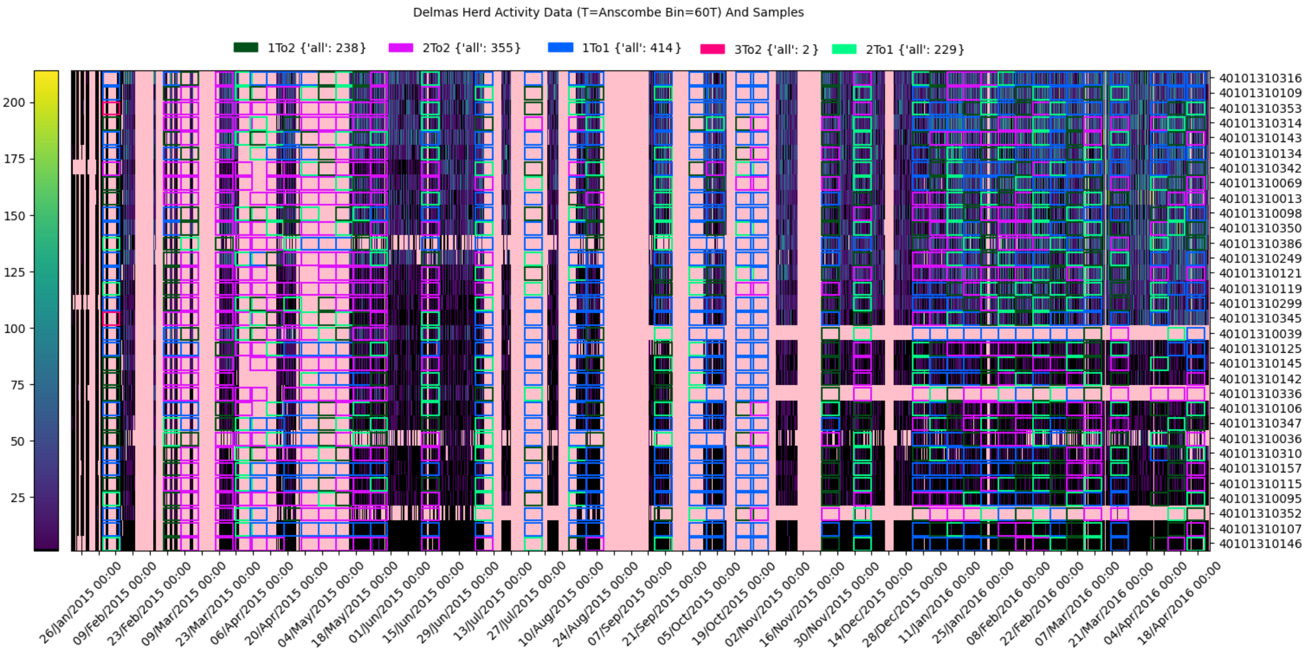


**B** – Goats

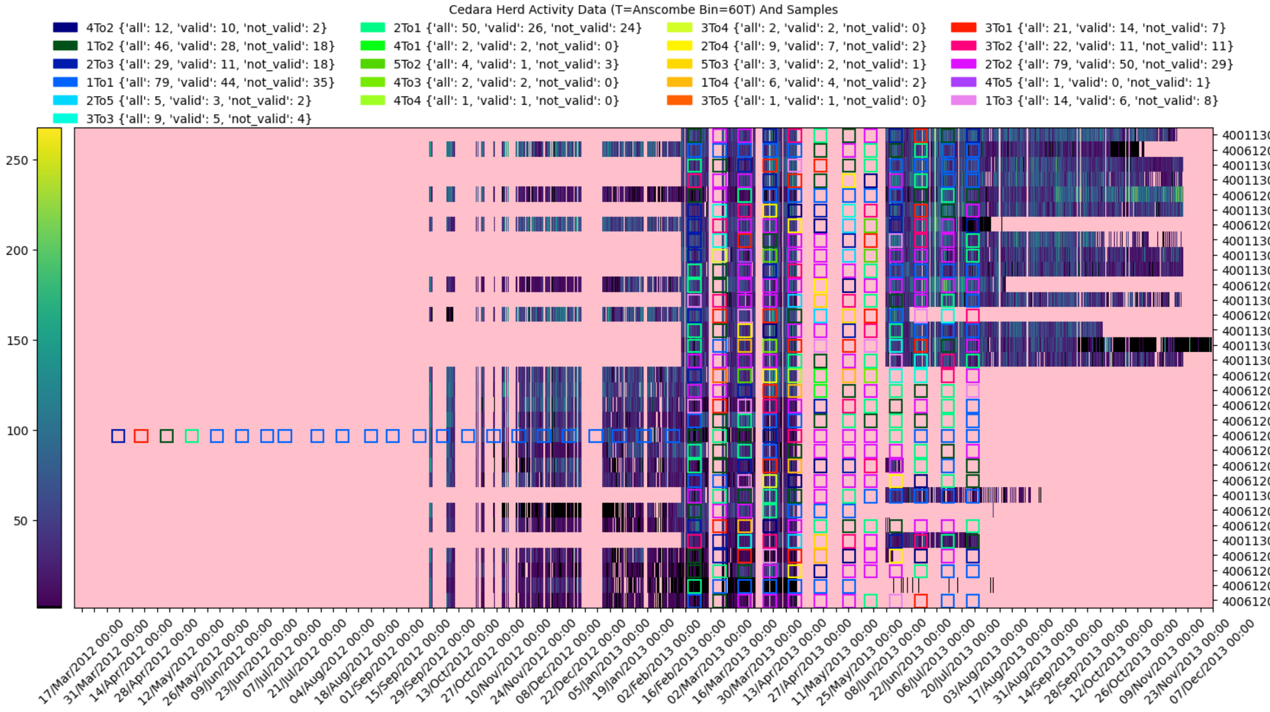


**Fig. S22. Heatmaps of FAMACHA samples for ML**. For both heatmaps the x-axis shows the time from the start to the end of the study, the y-axis shows the different transponders in the herd after filtering out invalid ones (Section. B.2). The activity count data displayed is transformed with the Log of the Anscombe (Sections. B.2, B.1). The data is re-sampled to an hour resolution (i.e. each bin contains the sum of 60 activity counts) for ease of visualisation of the entire time frame. While (A) show the sheep herd (B) shows the goat herd. The FAMACHA transitions are located and specified as per the legend with the overlay of coloured boxes. Note that in these heatmaps the activity data is not imputed, missing points are highlighted by the pink background.

Because we use supervised ML, we need to build a dataset which contains sets of features and target/label or ground truth, i.e. the value we want to predict. In our case, the label is the change of FAMACHA score from two consecutive tests, the time distance between two tests being a week for the sheep farm Delmas and two weeks for the goat farm Cedara. The features in each sample are composed of the activity time series associated with a FAMACHA transition. The feature time series is selected based on the date of the first FAMACHA test, it contains the activity data measured before the test date at midnight, for example for a FAMACHA transition from 1 to 2 with the first FAMACHA score evaluated at 2 on Feb 12, a feature array contains 1 day of activity data will contain the data measured from Feb 11 at 00:00am to Feb 12 at 00:00pm (Fig S23). The date of the FAMACHA tests of all samples for the sheep herd and the goat herd can be visualised in Figure S22.

Class imbalance is a common problem in ML (17), where the distribution of instances across different classes is highly imbalanced, with one or more classes having significantly fewer instances than the others. This can lead to biased predictions, where the model tends to favour the majority class and performs poorly on the minority class. There are various techniques and algorithms that can be used to handle class imbalance, such as resampling, cost-sensitive learning, and ensemble methods (18). For the FAMACHA transitions $\leq 2$ our dataset did not have a significant class imbalance between the healthy and unhealthy (1To2 or 2To2) group as illustrated in Table S1. The biggest imbalance was in the case of the Sheep farm when comparing the healthy samples (1To1) with the unhealthy samples (1To2) where we had 156 and 98 possible examples respectively, in this case, the healthy group was approximately 1.6 times larger than the unhealthy. Slightly imbalanced datasets, such as in our use case, may not necessarily be a major concern for ML algorithms. In such cases, the class imbalance is relatively small, and many algorithms can still perform well without any specific handling of the imbalance (19).

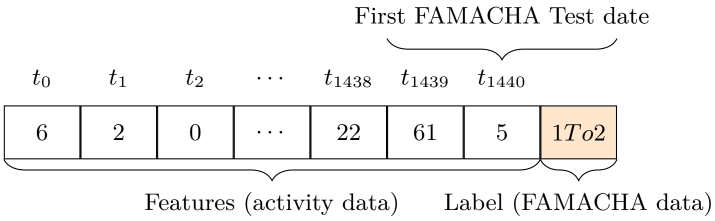


**Fig. S23. Example of a single day-long ML sample**. Illustration of a ML sample composed of 1440 features (all activity counts) and a FAMACHA label indicating a transition of FAMACHA from 1 to 2. Note that we only use activity data sampled at the 1-minute resolution, in this example there is a day (1440 minutes) of activity data, 1 minute per feature.

As this research focuses on a binary classification approach, we decided to build 2 groups of FAMACHA transitions, one being "healthy" and the other "unhealthy". Each group can contain single or multiple FAMACHA transitions. We tried groups of different compositions to find the optimal configuration for accurate classification. we explained that FAMACHA scores $\geq 3$ indicate poor health of the animal but our dataset is mostly composed of FAMACHA transitions containing score 1 and 2. We will define a sample as healthy if the FAMACHA score remained 1 for the two consecutive tests and we will refer to this transition with the notation FAMACHA $1 \rightarrow 1$ (1To1). Similarly we will refer to the unhealthy samples as FAMACHA $1 \rightarrow 2$ (1To2) or FAMACHA $2 \rightarrow 2$ (2To2). Although the amount of samples with other FAMACHA transitions is too low for training, we can test our models against those samples. Intuitively the unhealthy samples with FAMACHA $2 \rightarrow 2$ transition will be in a more "unhealthy" state that the FAMACHA $1 \rightarrow 2$ as the animal remained at a higher FAMACHA score (2) for longer, we will validate this hypothesis in the result section.

As part of routine husbandry, each animal which scored $\geq 2$ during a FAMACHA evaluation was immediately treated with Levamisole (Ripercol-L, Bayer Animal Health) at $7.5\,mg.kg^{-1}$ (20). We created a dataset to examine our ability to predict which animals responded well to treatment, comparing FAMACHA $2 \rightarrow 1$ against FAMACHA $1 \rightarrow 1$. This was based on classifying the 5 days of accelerometer data immediately following treatment for the sheep. For the goat, FAMACHA $2 \rightarrow 1$ against FAMACHA $1 \rightarrow 1$ was compared and 3 days of accelerometer data were used.

| | **Cedara** | **Delmas** |
|---|---|---|
| **Healthy** | 1To1 | 1To1 |
| **Unhealthy** | 2To2, 1To2 | 2To2, 1To2 |

**Table S1. Definition of healthy/unhealthy status**

**A. Intensity normalisation.** The magnitude or scale of the activity data greatly changes depending on the transponder used. The accelerometry sensor calibration might not be uniform across all of the transponders used. Furthermore, despite attaching the transponders with the same collar in the same position, it is challenging to regulate the tightness of the attachment. In addition, the attachment used does loosen up over time which will affect the recorded accelerometry values as the device will move with less inertia. For these reasons, intensity normalisation at the sample level is key for our ML approach as we aim to model the difference in patterns that might indicate ill-health within our data and avoid bias due to the difference in intensities that we have no control over.

Feature scaling is an important step for most ML algorithms, some algorithm such as K-NN depends on computing distances between the data points while others rely on solving the gradient descent optimisation problem which is highly dependent on the scale of each feature. If a feature has an order of magnitude larger than others, it might have an overly high weight in the objective function of the algorithm and make it unable to learn from other features correctly. Features can have completely different magnitude ranges naturally, especially if they come from different sources, for example, weather data such as temperature and rainfall have different magnitude ranges. Feature normalisation aims to scale all the features to a comparable scale of the same range.

Some of the commonly used techniques for feature scaling include L2-normalisation, Min-Max normalisation and Standard Scaling. Min-Max normalisation scales the data array $x$ between the range $[0, 1]$:

$$x_{scaled} = \frac{x - x_{min}}{x_{max} - x_{min}} \quad [8]$$

While Standard Scaling subtracts the input array $x$ by its mean value and divides it by its standard deviation $\sigma$, which centres the data around zero and scales it to unit variance ($\sigma = 1$).

$$x_{scaled} = \frac{x - \overline{x}}{\sigma} \quad [9]$$

L2 Normalisation simply divides the input sample array $x$ by its Euclidean distance:

$$||x||_2 = \sqrt{\sum_{i=1}^{n} x_i^2} = \sqrt{x_1^2 + x_2^2 + \cdots + x_n^2} \quad [10]$$

These techniques are all linear transformations which do not change the probability distribution of the data, they only scale the magnitude of the data. Each of these techniques could be used for sample normalisation, however, as all the data points are used to compute the scaling they can be biased by random bursts of high or low activity. Rather, we are interested in normalising each sample robustly so that only data points comparative to herd activity levels are taken into account. A viable solution is Quotient normalisation (QN), which scales the intensity of samples originating from each animal with a scaling factor that takes into consideration the herd behaviour.

### 4. Exogenous factors

We have discussed how the parasitic burden of small ruminants is directly linked to the life cycle of *Helminths*. We know that the worm population exponentially increase in wet weather, we can use the rainfall data on the location of each farm to asses how much predictive power it holds in comparison with the activity data, we will also combine the activity and rainfall data as well as other exogenous data sources such as the temperature and the wind speed. To include exogenous data in our samples we will append the data in the feature array (Fig S24). Similarly to the activity data (3), we select the data before the first FAMACHA test date up to the length of the sample. However, contrary to the activity data which is sampled per minute, the resolution (measurement per unit of time) of the weather data is hourly, it is not necessary to upscale the weather data to match the activity data resolution as this would only add redundant values to the feature array and the ML algorithms we used are not sensitive to the concept of different time resolutions in the feature space. We will only use Min Max scaling (section A) which preserves the relative differences in magnitude while bringing all values within a standardised interval. This approach preserves the original scale of the data while still providing a consistent range for machine learning algorithms that are sensitive to feature scales such as SVM.

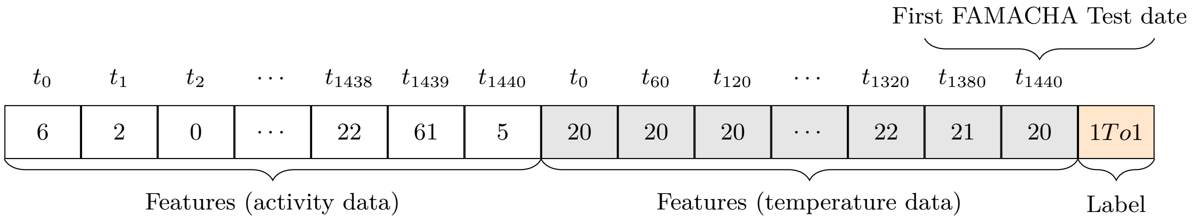


**Fig. S24. Example of a single sample with exogenous data**. Illustration of a ML sample composed of activity and temperature data and a FAMACHA label indicating a transition of FAMACHA from 1 to 1. Note that the temperature data is sampled hourly while the activity data is sampled per minute.

**A. Adding exogenous features.** In figures S27 we added exogenous features in our samples or only used exogenous features without activity data (Fig S24). The parasitic burden of the animals is directly correlated to the rainfall on the farm. As expected when using the rainfall data (black box plot) the predictive power of our SVM model was high with a 76% median AUC with the 84 days of rainfall data before the FAMCHA test, the predictive power reduced significantly to 65% AUC when

**A** – Goats

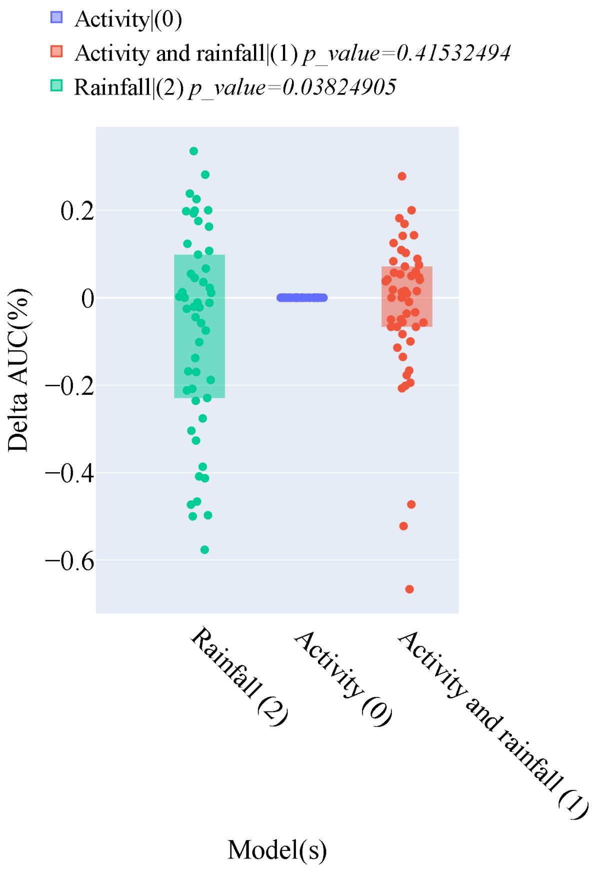


**Fig. S25. Performance of SVM model trained on activity against Rainfall or both.** The left y-axis shows the number of samples while the right axis shows the difference compared to the best model AUC.

using a short 7 days window of the rainfall before the test which can be explained by the fact that the worm hatching time onset. Using a mix of Activity data and rainfall (orange box plot) gave an AUC above 80%.

While rainfall is known to have a direct link to the life cycle of *H. contortus*, we also studied the impact of other exogenous factors with the same method, namely ambient air humidity, temperature and windspeed in S26 where we compared the predictive power of our model trained with activity data against each of those factors or a combination of activity and exogenous feature. While the results are inconclusive for the goat farm we found that on the Sheep farm temperature alone had similar predictive power as rainfall although activity remained the best predictor.

**A** – Sheep

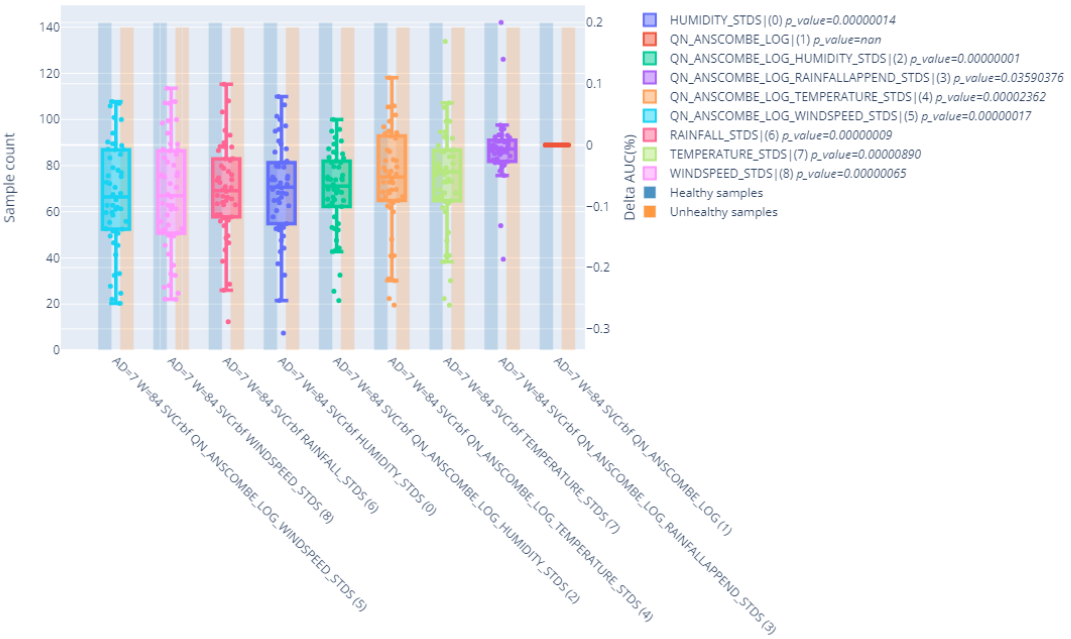


**B** – Goat

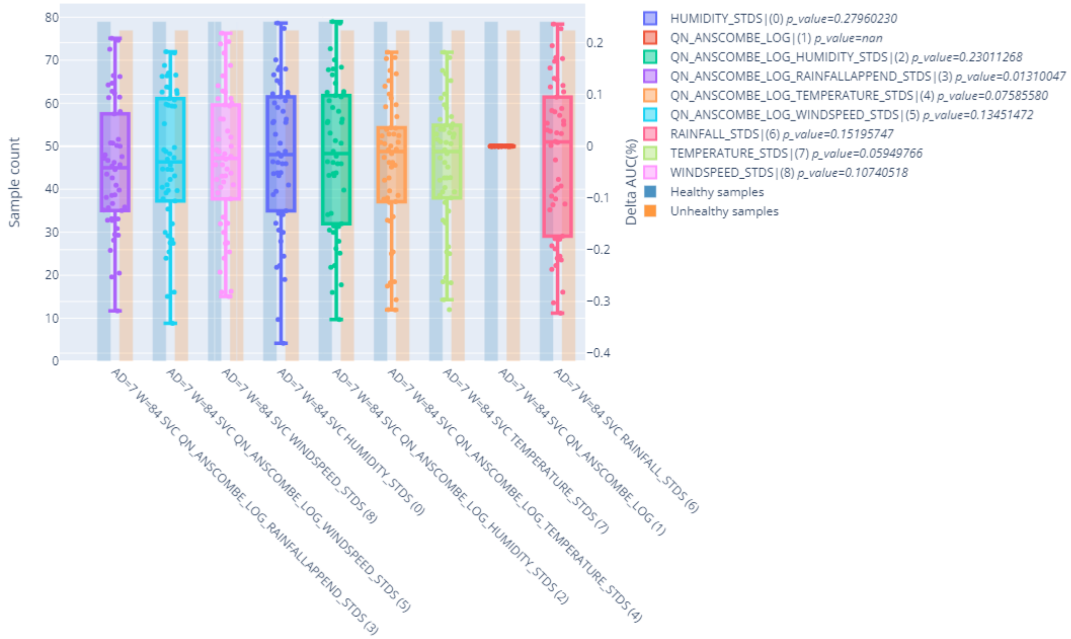


**Fig. S26. Performance of SVM model with exogenous features compared to model trained with activity only.** The models trained with a combination of activity and rainfall data are annotated with the word "APPEND" in the legend. The left y-axis shows the number of samples while the right axis shows the difference compared to the best model AUC.

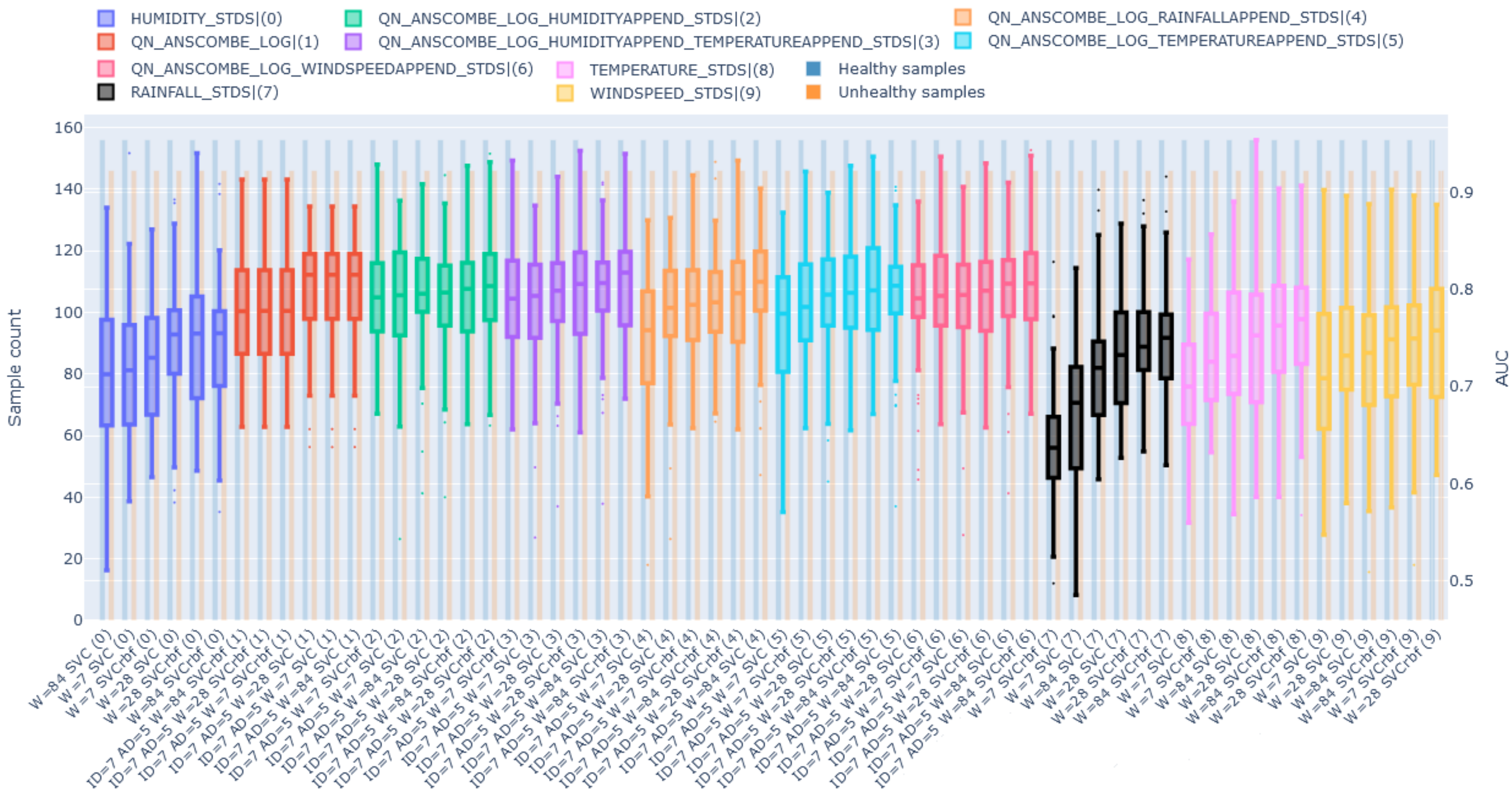


**Fig. S27. Performance of SVM model with exogenous features for the sheep farm.** The x-axis shows the number of activity days via the notation "AD=", the number of synthetic days with "ID=" and "W=" for the number of days for the weather. For example, if the x-axis shows "ID=7 AD=5 W=84 SVC" this means that if the samples contain activity data, a sample can contain up to 7 synthetic days, the activity time series contains 5 days from the first famacha test and the sample contains 84 days of weather (rainfall, temperature or wind speed) data. The models trained with a combination of activity and weather features are annotated with the word "APPEND" in the legend. The left y-axis shows the number of samples while the right axis shows the AUC.

### 5. Classification of health status

For both farms, we will compare the results obtained by training different types of models, namely SVM (linear and rbf kernel), logistic regression (LREG), decision tree (DTREE) and k-nearest-neighbours (KNN). We will also use different training labels for the health status of the samples, FAMACHA score of $1 \rightarrow 1$ will be the healthy class while FAMACHA score of $2 \rightarrow 2$ or FAMACHA score of $1 \rightarrow 2$ will be unhealthy. The number of activity days in a sample will be changed to 1, 4 or 7. Furthermore, we will apply different pre-processing approaches to the activity time series of the samples. Initially we will show the top and bottom test results based on the AUC performance of the different models in tables S2 and S3. In figures S34 and S35 we show that the predictive power of our model is higher when we train with unhealthy samples with FAMACHA score of $2 \rightarrow 2$ compared to $1 \rightarrow 2$, this is likely due to the animal being in an unhealthy state (FAMACHA 2) for longer. Among the ML algorithms we tried SVM and LREG performed best with similar results while KNN and DTREE had lower performance. (Fig S31). In addition, although more computationally expensive we found that using the frequency domain CWT transform did not improve classification performance (Table S4).

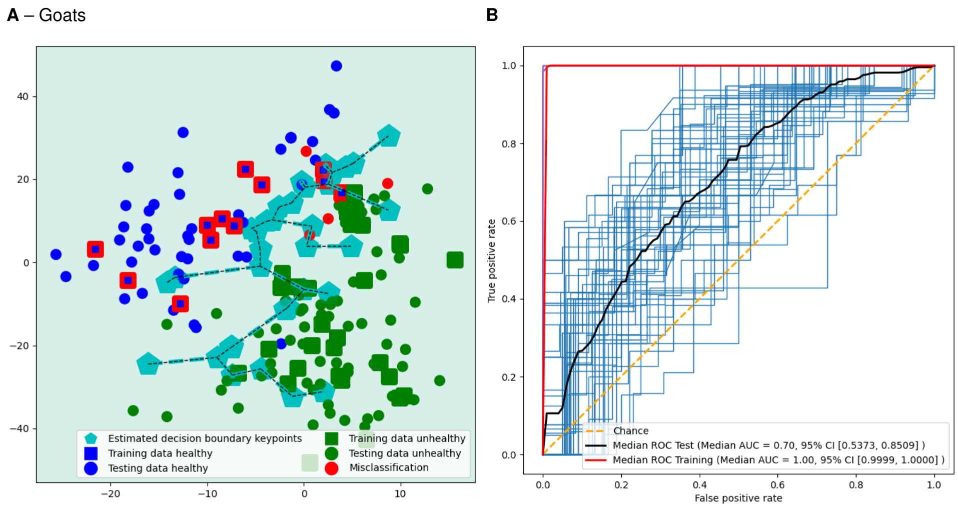


**Fig. S28. Classifying health status.** On the goat farm the machine learning was trained to discriminate FAMACHA $1 \rightarrow 1$ against FAMACHA $2 \rightarrow 2$ using 7 days of accelerometry data directly preceding the FAMACHA evaluation. (A) Decision boundary plot and (B) Receiver Operating Characteristic (ROC) curve for training and testing on goats.

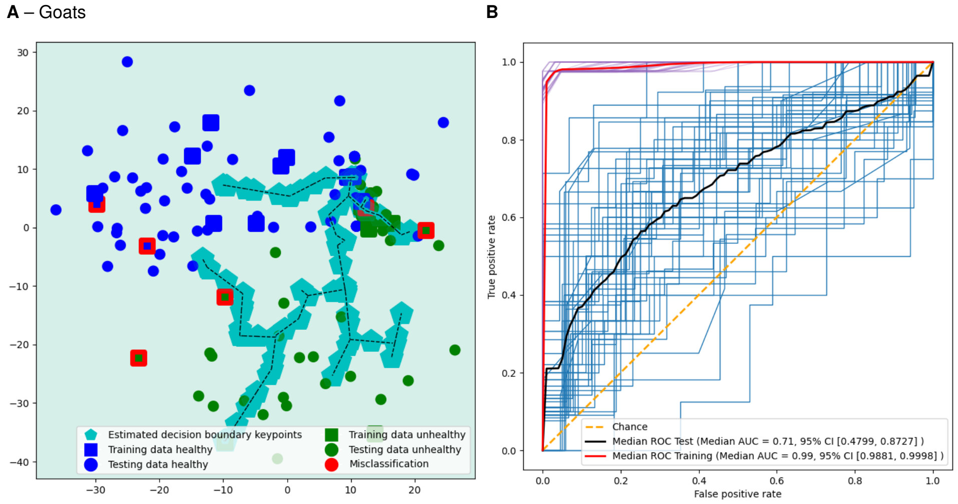


**Fig. S29. Classifying the response to treatment.** For both farms, the machine learning was trained to discriminate FAMACHA $2 \rightarrow 1$ against FAMACHA $1 \rightarrow 1$ using 2 days of accelerometry data directly preceding anthelmintic treatment for the Goat farm. (A) Decision boundary plot and (B) Receiver Operating Characteristic (ROC) curve for training and testing on goats.

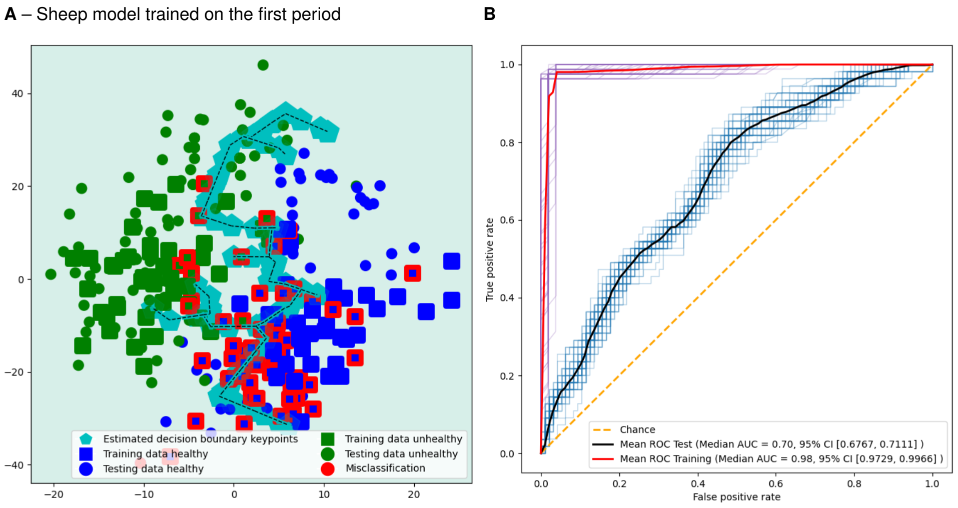


**Fig. S30. Temporal validation.** On both farms we used 6 days of accelerometry data directly preceding FAMACHA evaluation. On the sheep farm, the machine learning was trained to discriminate FAMACHA 1 → 1 against FAMACHA 2 → 2. To assess concept drift the dataset was split to obtain two equally sized consecutive periods. (A) Scatter plot and (B) ROC curve for training on the first period and testing on the second.

**A** – Sheep farm | Models trained with **unhealthy label 1To2**.

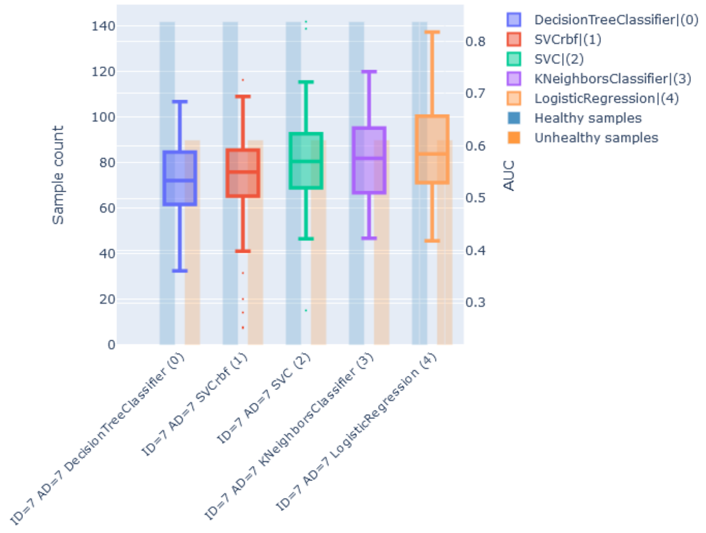


**B** – Sheep farm | Models trained with **unhealthy label 2To2**.

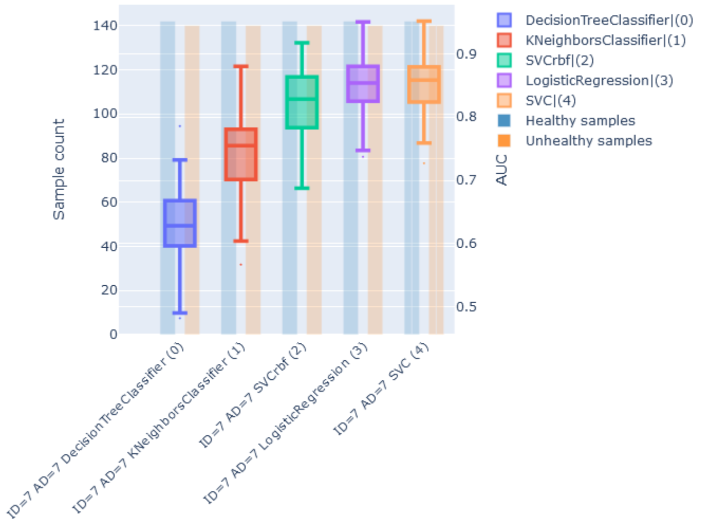


**C** – Goat farm | Models trained with **unhealthy label 1To2**.

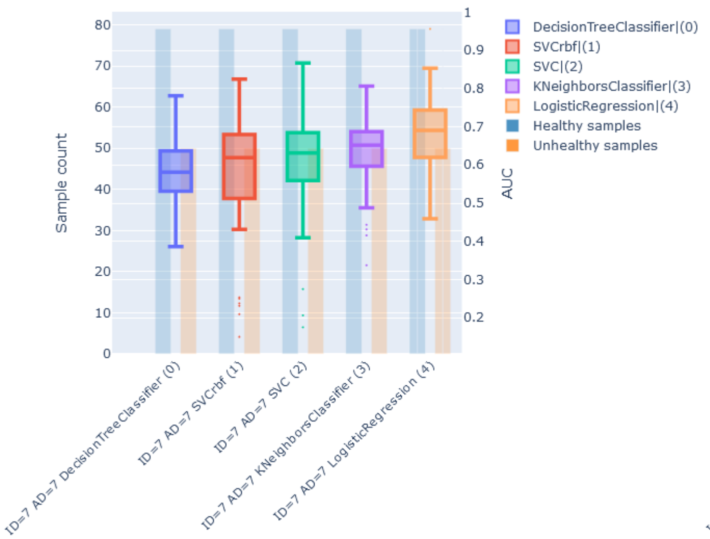


**D** – Goat farm | Models trained with **unhealthy label 2To2**.

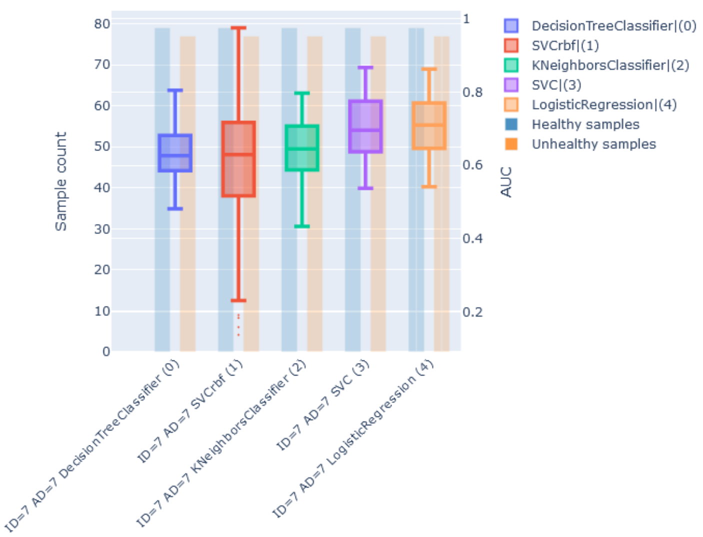


**Fig. S31. Performance of different ML algorithms with M-RNN imputed samples for different unhealthy labels.** By fixing the number of activity days and the maximum number of synthetic days allowed in the samples to 7, we can compare the performances of different type of models. We also fixed the pre-processing pipeline of the activity time series to QN → ANSCOMBE → LOG → STDS. For each figure, the x-axis shows the ML algorithm while the left y-axis shows the number of samples per class and the right y-axis shows the AUC.

| AUC test(95% CI) | Class0 P-test | Class1 P-test | N test | A-days | S-days | Imp | Clf | Pre-proc | Class1 | Farm |
|---|---|---|---|---|---|---|---|---|---|---|
| 0.80 (0.68-0.91) | 0.70 (0.53-0.92) | 0.72 (0.61-0.85) | 42 | 7 | 4 | M-RNN | LREG | QN->ANSCOMBE->LOG | 2To2 | delmas |
| 0.80 (0.67-0.91) | 0.70 (0.54-0.85) | 0.73 (0.62-0.89) | 42 | 7 | 4 | M-RNN | SVM(linear) | QN->ANSCOMBE->LOG | 2To2 | delmas |
| 0.80 (0.67-0.90) | 0.70 (0.53-0.85) | 0.73 (0.62-0.86) | 42 | 7 | 4 | M-RNN | SVM(linear) | QN->ANSCOMBE->LOG->MINMAX | 2To2 | delmas |
| 0.80 (0.67-0.90) | 0.70 (0.54-0.90) | 0.72 (0.60-0.85) | 42 | 7 | 4 | M-RNN | LREG | QN->ANSCOMBE->LOG->MINMAX | 2To2 | delmas |
| 0.80 (0.68-0.90) | 0.71 (0.53-0.85) | 0.74 (0.63-0.89) | 42 | 7 | 4 | M-RNN | SVM(linear) | QN->ANSCOMBE->LOG->STD | 2To2 | delmas |
| 0.80 (0.68-0.90) | 0.71 (0.53-0.85) | 0.74 (0.63-0.89) | 42 | 7 | 4 | M-RNN | SVM(linear) | | 2To2 | delmas |
| 0.80 (0.67-0.90) | 0.70 (0.53-0.84) | 0.73 (0.63-0.86) | 42 | 7 | 4 | M-RNN | LREG | | 2To2 | delmas |
| 0.80 (0.67-0.90) | 0.70 (0.53-0.84) | 0.73 (0.63-0.86) | 42 | 7 | 4 | M-RNN | LREG | QN->ANSCOMBE->LOG->STD | 2To2 | delmas |
| 0.77 (0.60-0.91) | 0.71 (0.50-0.86) | 0.69 (0.58-0.83) | 42 | 7 | 4 | M-RNN | LREG | QN | 2To2 | delmas |
| 0.77 (0.62-0.89) | 0.69 (0.53-0.85) | 0.70 (0.58-0.82) | 42 | 7 | 4 | M-RNN | SVM(linear) | QN | 2To2 | delmas |

**(a) Top 10 models.**

| AUC test(95% CI) | Class0 P-test | Class1 P-test | N test | A-days | S-days | Imp | Clf | Pre-proc | Class1 | Farm |
|---|---|---|---|---|---|---|---|---|---|---|
| 0.46 (0.31-0.61) | 0.61 (0.57-0.64) | 0.19 (0.00-1.00) | 41 | 7 | 4 | LI | SVM(linear) | L2 | 1To2 | delmas |
| 0.46 (0.30-0.64) | 0.60 (0.59-0.61) | 0.00 (0.00-0.00) | 33 | 4 | 1 | LI | SVM(rbf) | L2 | 1To2 | delmas |
| 0.45 (0.26-0.66) | 0.54 (0.42-0.63) | 0.39 (0.14-0.73) | 26 | 4 | 1 | M-RNN | LREG | QN | 1To2 | delmas |
| 0.45 (0.32-0.61) | 0.60 (0.59-0.61) | 0.00 (0.00-0.00) | 33 | 7 | 1 | LI | SVM(rbf) | L2 | 1To2 | delmas |
| 0.45 (0.32-0.56) | 0.61 (0.59-0.62) | 0.02 (0.00-0.31) | 41 | 7 | 4 | LI | SVM(rbf) | QN->ANSCOMBE->LOG->STD | 1To2 | delmas |
| 0.45 (0.32-0.56) | 0.61 (0.59-0.62) | 0.02 (0.00-0.31) | 41 | 7 | 4 | LI | SVM(rbf) | | 1To2 | delmas |
| 0.45 (0.28-0.64) | 0.57 (0.56-0.58) | 0.00 (0.00-0.00) | 26 | 4 | 1 | M-RNN | LREG | L2 | 1To2 | delmas |
| 0.44 (0.32-0.59) | 0.55 (0.52-0.57) | 0.04 (0.00-0.77) | 34 | 7 | 4 | M-RNN | LREG | L2 | 1To2 | delmas |
| 0.43 (0.29-0.55) | 0.61 (0.60-0.62) | 0.00 (0.00-0.00) | 41 | 7 | 4 | LI | SVM(rbf) | QN | 1To2 | delmas |
| 0.43 (0.24-0.62) | 0.57 (0.56-0.58) | 0.00 (0.00-0.00) | 26 | 7 | 1 | M-RNN | LREG | L2 | 1To2 | delmas |

**(b) Bottom 10 models.**

**Table S2. Comparison of model results for FAMACHA samples classification on the sheep farm. "Class0" and "Class1" refer to the healthy FAMACHA transition ("1To1") and the unhealthy transition. "P" refer to the model precision. "A" and "S" indicate "Activity" and "Synthetic" respectively. "Imp" shows the imputation technique used. "Clf" shows the type of model used. "Pre-proc" shows the preprocessing used on the sample time series before training (QN: Quotient Normalisation, "STD": Standard Scaling).**

| AUC test(95% CI) | Class0 P-test | Class1 P-test | N test | A-days | S-days | Imp | Clf | Pre-proc | Class1 | Farm |
|---|---|---|---|---|---|---|---|---|---|---|
| 0.68 (0.39-0.94) | 0.60 (0.50-0.78) | 0.69 (0.27-1.00) | 14 | 4 | 1 | LI | LREG | L2 | 2To2 | cedara |
| 0.66 (0.45-0.89) | 0.62 (0.41-0.84) | 0.61 (0.34-0.97) | 14 | 4 | 1 | M-RNN | LREG | QN->ANSCOMBE->LOG | 2To2 | cedara |
| 0.66 (0.45-0.88) | 0.61 (0.43-0.83) | 0.60 (0.34-0.96) | 14 | 4 | 1 | M-RNN | LREG | | 2To2 | cedara |
| 0.66 (0.45-0.88) | 0.61 (0.43-0.83) | 0.60 (0.34-0.96) | 14 | 4 | 1 | M-RNN | LREG | QN->ANSCOMBE->LOG->STD | 2To2 | cedara |
| 0.66 (0.45-0.88) | 0.62 (0.41-0.84) | 0.61 (0.38-0.96) | 14 | 4 | 1 | M-RNN | LREG | QN->ANSCOMBE->LOG->MINMAX | 2To2 | cedara |
| 0.64 (0.41-0.87) | 0.56 (0.47-0.67) | 0.69 (0.00-1.00) | 14 | 4 | 1 | M-RNN | LREG | L2 | 2To2 | cedara |
| 0.64 (0.31-0.88) | 0.57 (0.43-0.70) | 0.65 (0.00-1.00) | 14 | 7 | 1 | LI | LREG | L2 | 2To2 | cedara |
| 0.64 (0.31-0.89) | 0.66 (0.00-1.00) | 0.57 (0.39-0.81) | 14 | 4 | 1 | LI | SVM(linear) | QN | 2To2 | cedara |
| 0.64 (0.39-0.91) | 0.68 (0.08-1.00) | 0.57 (0.40-0.82) | 14 | 4 | 1 | LI | LREG | QN | 2To2 | cedara |
| 0.64 (0.17-0.87) | 0.62 (0.41-0.84) | 0.61 (0.35-0.97) | 14 | 4 | 1 | M-RNN | SVM(linear) | QN->ANSCOMBE->LOG | 2To2 | cedara |

**(a) Top 10 models.**

| AUC test(95% CI) | Class0 P-test | Class1 P-test | N test | A-days | S-days | Imp | Clf | Pre-proc | Class1 | Farm |
|---|---|---|---|---|---|---|---|---|---|---|
| 0.42 (0.10-0.77) | 0.54 (0.42-0.62) | 0.51 (0.00-1.00) | 14 | 4 | 1 | M-RNN | SVM(rbf) | QN->ANSCOMBE->LOG->STD | 2To2 | cedara |
| 0.41 (0.20-0.64) | 0.68 (0.55-0.82) | 0.44 (0.00-1.00) | 14 | 7 | 4 | LI | SVM(linear) | QN->ANSCOMBE->LOG->MINMAX | 1To2 | cedara |
| 0.41 (0.10-0.75) | 0.55 (0.44-0.62) | 0.66 (0.00-1.00) | 14 | 4 | 1 | M-RNN | SVM(rbf) | QN->ANSCOMBE->LOG | 2To2 | cedara |
| 0.41 (0.15-0.77) | 0.56 (0.47-0.63) | 0.68 (0.00-1.00) | 14 | 7 | 1 | LI | SVM(rbf) | L2 | 2To2 | cedara |
| 0.41 (0.18-0.66) | 0.56 (0.46-0.67) | 0.62 (0.00-1.00) | 14 | 7 | 1 | M-RNN | SVM(linear) | L2 | 2To2 | cedara |
| 0.41 (0.13-0.76) | 0.53 (0.50-0.57) | 0.22 (0.00-1.00) | 14 | 4 | 1 | M-RNN | SVM(rbf) | L2 | 2To2 | cedara |
| 0.41 (0.10-0.74) | 0.54 (0.42-0.62) | 0.56 (0.00-1.00) | 14 | 4 | 1 | M-RNN | SVM(rbf) | QN->ANSCOMBE->LOG->MINMAX | 2To2 | cedara |
| 0.40 (0.08-0.71) | 0.56 (0.45-0.67) | 0.67 (0.08-1.00) | 14 | 7 | 1 | LI | SVM(rbf) | | 2To2 | cedara |
| 0.40 (0.08-0.71) | 0.56 (0.45-0.67) | 0.67 (0.08-1.00) | 14 | 7 | 1 | LI | SVM(rbf) | QN->ANSCOMBE->LOG->STD | 2To2 | cedara |
| 0.40 (0.12-0.74) | 0.56 (0.46-0.62) | 0.72 (0.08-1.00) | 14 | 7 | 1 | LI | SVM(rbf) | QN->ANSCOMBE->LOG | 2To2 | cedara |

**(b) Bottom 10 models.**

**Table S3. Comparison of model results for FAMACHA samples classification on the goat farm. "Class0" and "Class1" refer to the healthy FAMACHA transition ("1To1") and the unhealthy transition. "P" refer to the model precision. "A" and "S" indicate "Activity" and "Synthetic" respectively. "Imp" shows the imputation technique used. "Clf" shows the type of model used. "Pre-proc" shows the preprocessing used on the sample time series before training (QN: Quotient Normalisation, "STD": Standard Scaling).**

The training AUC (Area Under the Curve) is a measure of how well a ML model can distinguish between positive and negative classes in the training dataset. It represents the performance of the model on data that it has already seen during the training process. The testing AUC, on the other hand, measures the model's performance on a completely independent dataset that it has not seen during training. It gives an estimate of how well the model will perform on new, unseen data. If the training AUC is higher than the testing AUC, it indicates that the model is overfitting to the training data. Overfitting occurs when a model learns to capture noise and random fluctuations in the training data, rather than the underlying patterns that generalise to new data. To address overfitting, we can use techniques such as regularisation, in practice, we would have to tune the parameter of our models with a parameter grid search which is timely and computationally expensive. In figure S32A we show the training and testing AUCs of one of our LREG model and in figure S32B one of our SVM models, we can observe that although both models overfit on the training data with the default regularisation parameters the still performed well on the testing datasets, fine-tune parameter tuning may increase the testing performances even further.

**A** – Sheep farm | Models trained with **unhealthy label 2To2** while healthy is 1To1. **B** – Goat farm | Models trained with **unhealthy label 2To2** while healthy is 1To1.

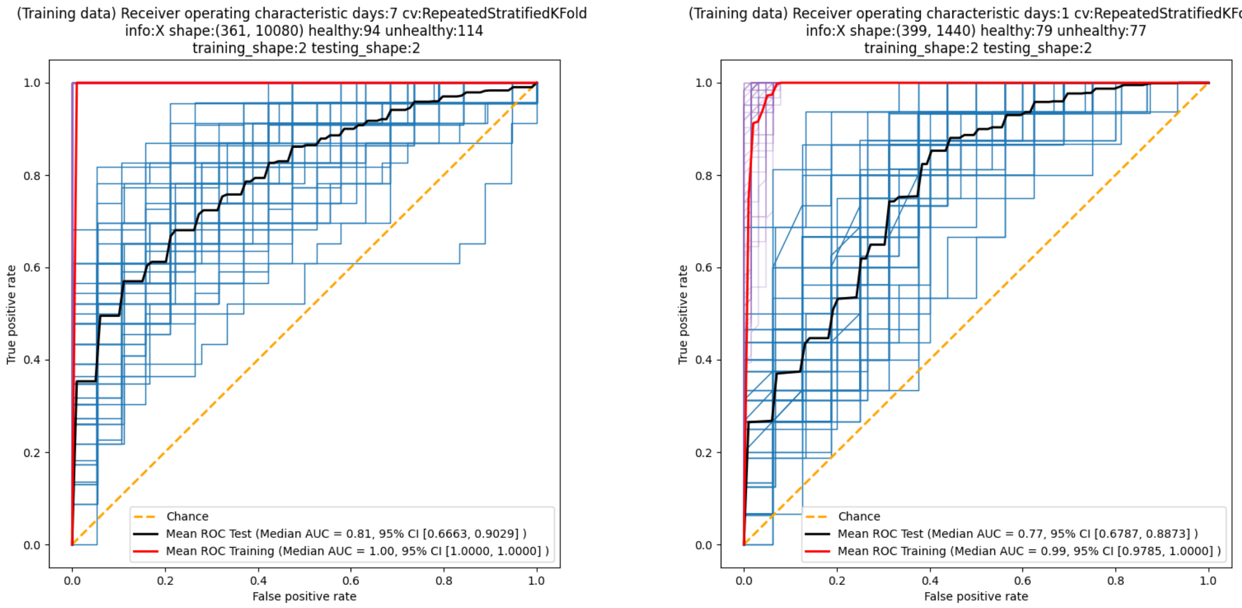


**Fig. S32. ROC curves of models for classification of health status with Logistic regression model.** For each figure, the black curve shows the median test auc from the cross-validation while the blue curves show the AUC of each individual test split. Similarly, the red curve shows the median AUC for the training data while the purple curves show the AUC of each training split.

**A. Model evaluation on unseen labels.** We plot density histograms of the prediction probability of our cross-validated best model (Tables. S3a and S2a) to evaluate how well the model classifies unseen samples. The model will classify a sample as unhealthy if the probability output is $\geq 0.5$ (marked as a vertical grey dotted line) (Fig S36).

We show the result for the sheep farm in figure S33A. An SVM model was trained on healthy famacha (1To1, 142 examples) and unhealthy FAMACHA (2To2, 140 examples) and tested with repeated k-fold cross-validation on all of the samples available in the dataset including the ones used for training. When the density histogram values are skewed to the left it means that the model classifies the data as healthy while on the opposite side, it will classify as unhealthy. As expected the model perform well by properly classifying the label used for training "1To1" and "2To2" while the "1To2" and "2To1" transitions were mostly classified as healthy, arguably both are still in the healthy range of FAMACHA.

The goat farm figure S36A shows the results. Similarly, An SVM model was trained on healthy famacha (1To1, 79 examples) and unhealthy FAMACHA (2To2, 79 examples). Although in small numbers the goat farm had more variation of FAMCHA transitions that our model could be tested on, as expected the density histogram on the trained label shows mostly correct classifications, for the advanced stage in FAMCHA such as "2To5", "4To5", "5To3", "5To2", "4To3" and "4To1" our model perfectly classified all samples as unhealthy, even for the "1To2" transition the model seems to start "understanding" health decline as a majority of samples were classified as unhealthy instead of healthy with the demarcation getting clearer with "1To3", "1To4", "2To3", "2To4", "3To4", "3To1", "3To2" and "4To2". Only "3To5" showed mitigated results where the model performance was poor.

**A** – Sheep farm | SVM(rbf) trained with **unhealthy label 1To2** while healthy is 1To1

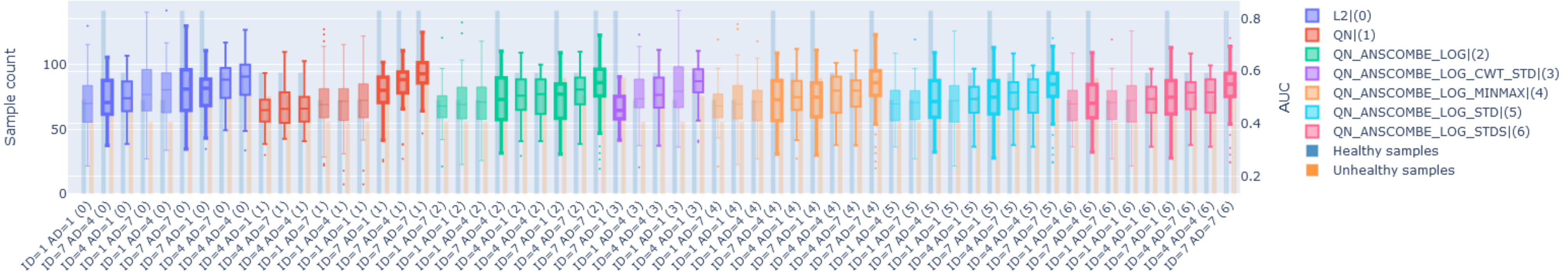


**B** – Sheep farm | SVM(rbf) trained with **unhealthy label 2To2** while healthy is 1To1

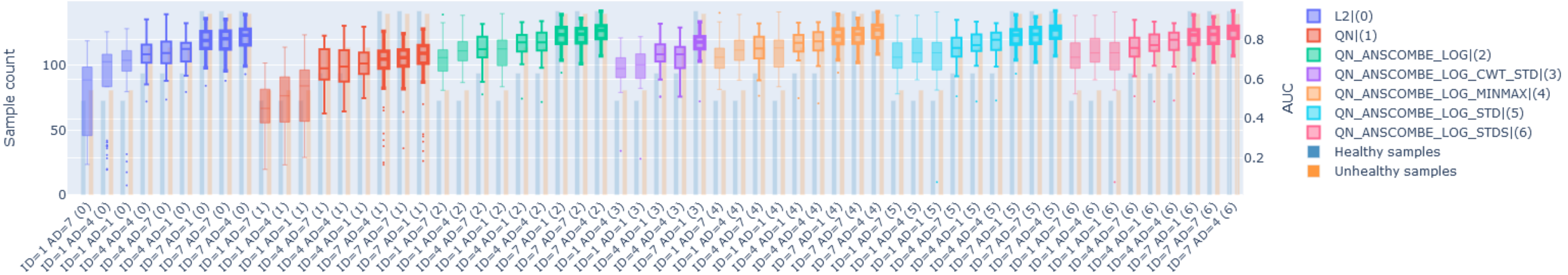


**Fig. S34. Performance of SVM(rbf) with M-RNN imputed samples for different unhealthy labels on the sheep farm.** Boxplot illustration of different SVM models trained with different versions of the samples. The x-axis shows the number of activity days used in the samples with the notation "AD=", for example, "AD=1" means that the samples contained 1 day of activity data, similarly "ID=" indicates the number of fully synthetic(imputed) days allowed in the sample, the boxplot line thickness also shows the amount of imputed data allowed in the samples. The preprocessing pipeline is indicated by the integer in parenthesis, for example (0) refers to applying L2 normalisation on the samples' time series. The y-axis on the left shows the number of samples in the healthy and unhealthy classes while the right axis shows the AUC.

**A** – Goat farm | SVM(rbf) trained with **unhealthy label 1To2** while healthy is 1To1

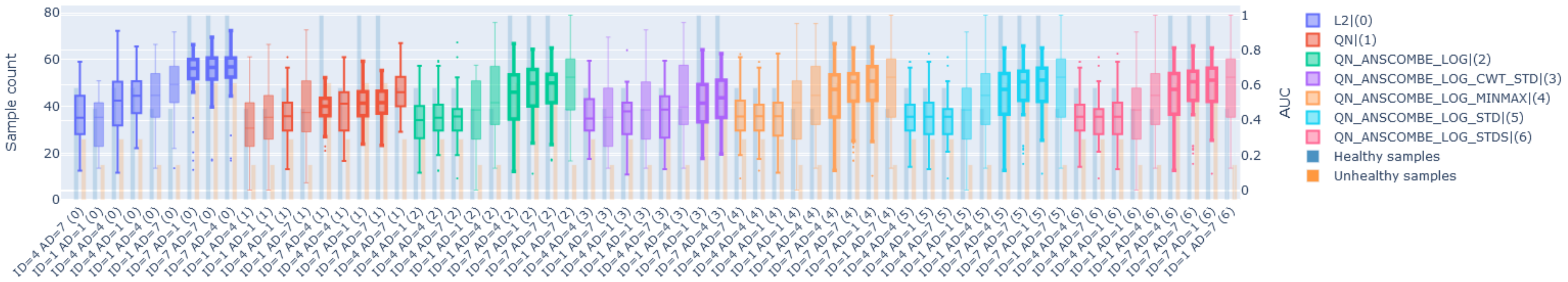


**B** – Goat farm | SVM(rbf) trained with **unhealthy label 2To2** while healthy is 1To1

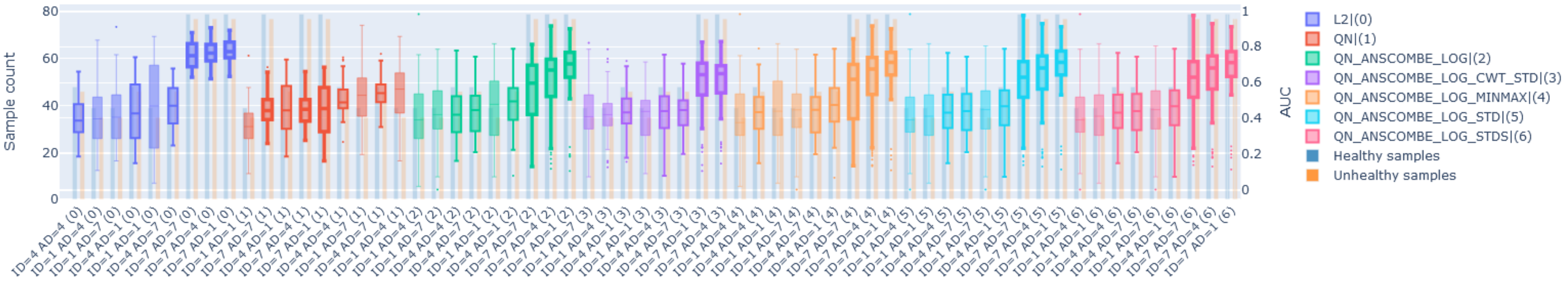


**Fig. S35. Performance of SVM(rbf) with M-RNN imputed samples for different unhealthy labels on the goat farm.** Boxplot illustration of different SVM models trained with different versions of the samples. The x-axis shows the number of activity days used in the samples with the notation "AD=", for example, "AD=1" means that the samples contained 1 day of activity data, similarly "ID=" indicates the number of fully synthetic(imputed) days allowed in the sample, the boxplot line thickness also shows the amount of imputed data allowed in the samples. The preprocessing pipeline is indicated by the integer in parenthesis, for example (0) refers to applying L2 normalisation on the samples' time series. The y-axis on the left shows the number of samples in the healthy and unhealthy classes while the right axis shows the AUC.

| AUC test(95% CI) | Class0 P-test | Class1 P-test | N test | A-days | S-days | Imp | Clf | Pre-proc | Class1 | Farm |
|---|---|---|---|---|---|---|---|---|---|---|
| 0.80 (0.80-0.80) | 0.83 (0.83-0.83) | 0.74 (0.74-0.74) | 31 | 7 | 1 | M-RNN | SVM(linear) | QN->ANSCOMBE->LOG->CWT->STD | 2To2 | delmas |
| 0.69 (0.58-0.81) | 0.63 (0.44-0.86) | 0.63 (0.50-0.78) | 31 | 7 | 1 | M-RNN | SVM(rbf) | QN->ANSCOMBE->LOG->CWT->STD | 2To2 | delmas |
| 0.65 (0.54-0.81) | 0.61 (0.43-0.82) | 0.63 (0.52-0.78) | 31 | 7 | 1 | M-RNN | LREG | QN->ANSCOMBE->LOG->CWT->STD | 2To2 | delmas |
| 0.65 (0.53-0.78) | 0.63 (0.46-0.87) | 0.62 (0.54-0.69) | 31 | 4 | 1 | M-RNN | SVM(rbf) | QN->ANSCOMBE->LOG->CWT->STD | 2To2 | delmas |
| 0.63 (0.44-0.80) | 0.67 (0.46-0.92) | 0.54 (0.44-0.66) | 37 | 7 | 1 | LI | KNN | QN->ANSCOMBE->LOG->CWT->STD | 2To2 | delmas |
| 0.61 (0.48-0.80) | 0.70 (0.41-1.00) | 0.57 (0.50-0.64) | 31 | 4 | 1 | M-RNN | KNN | QN->ANSCOMBE->LOG->CWT->STD | 2To2 | delmas |
| 0.61 (0.45-0.75) | 0.64 (0.43-0.87) | 0.53 (0.40-0.63) | 37 | 4 | 1 | LI | KNN | QN->ANSCOMBE->LOG->CWT->STD | 2To2 | delmas |
| 0.58 (0.44-0.71) | 0.55 (0.36-0.72) | 0.57 (0.47-0.69) | 31 | 4 | 1 | M-RNN | LREG | QN->ANSCOMBE->LOG->CWT->STD | 2To2 | delmas |
| 0.58 (0.42-0.76) | 0.61 (0.46-0.76) | 0.55 (0.39-0.79) | 37 | 7 | 1 | LI | DTREE | QN->ANSCOMBE->LOG->CWT->STD | 2To2 | delmas |
| 0.57 (0.39-0.71) | 0.60 (0.42-0.73) | 0.55 (0.36-0.75) | 37 | 4 | 1 | LI | DTREE | QN->ANSCOMBE->LOG->CWT->STD | 2To2 | delmas |

**(a) Sheep farm| Top 10 models with CWT.**

| AUC test(95% CI) | Class0 P-test | Class1 P-test | N test | A-days | S-days | Imp | Clf | Pre-proc | Class1 | Farm |
|---|---|---|---|---|---|---|---|---|---|---|
| 0.63 (0.29-0.89) | 0.73 (0.60-0.87) | 0.63 (0.00-1.00) | 14 | 7 | 4 | M-RNN | LREG | QN->ANSCOMBE->LOG->CWT->STD | 1To2 | cedara |
| 0.59 (0.33-0.86) | 0.55 (0.37-0.77) | 0.53 (0.33-0.75) | 18 | 7 | 4 | LI | LREG | QN->ANSCOMBE->LOG->CWT->STD | 2To2 | cedara |
| 0.59 (0.38-0.85) | 0.55 (0.42-0.70) | 0.54 (0.33-0.74) | 18 | 7 | 4 | M-RNN | LREG | QN->ANSCOMBE->LOG->CWT->STD | 2To2 | cedara |
| 0.59 (0.27-0.82) | 0.57 (0.40-0.82) | 0.51 (0.25-0.71) | 14 | 7 | 1 | M-RNN | LREG | QN->ANSCOMBE->LOG->CWT->STD | 2To2 | cedara |
| 0.58 (0.32-0.78) | 0.59 (0.33-0.78) | 0.58 (0.26-0.84) | 18 | 7 | 4 | M-RNN | DTREE | QN->ANSCOMBE->LOG->CWT->STD | 2To2 | cedara |
| 0.58 (0.25-0.83) | 0.69 (0.43-0.98) | 0.38 (0.03-0.61) | 14 | 7 | 4 | LI | KNN | QN->ANSCOMBE->LOG->CWT->STD | 1To2 | cedara |
| 0.57 (0.29-0.82) | 0.58 (0.27-0.81) | 0.52 (0.17-0.70) | 14 | 4 | 1 | LI | LREG | QN->ANSCOMBE->LOG->CWT->STD | 2To2 | cedara |
| 0.57 (0.27-0.86) | 0.59 (0.28-0.83) | 0.51 (0.22-0.71) | 14 | 7 | 1 | LI | LREG | QN->ANSCOMBE->LOG->CWT->STD | 2To2 | cedara |
| 0.56 (0.14-0.86) | 0.73 (0.57-0.88) | 0.63 (0.00-1.00) | 14 | 7 | 4 | M-RNN | SVM(linear) | QN->ANSCOMBE->LOG->CWT->STD | 1To2 | cedara |
| 0.55 (0.32-0.74) | 0.58 (0.31-0.83) | 0.52 (0.21-0.74) | 14 | 7 | 1 | LI | DTREE | QN->ANSCOMBE->LOG->CWT->STD | 2To2 | cedara |

**(b) Goat farm| Top 10 models with CWT.**

**Table S4. Comparison of model results for health classification with frequency domain data (CWT). "Class0" and "Class1" refer to the healthy FAMACHA transition ("1To1") and the unhealthy transition. "P" refer to the model precision. "A" and "S" indicate "Activity" and "Synthetic" respectively. "Imp" shows the imputation technique used. "Clf" shows the type of model used. "Pre-proc" shows the preprocessing used on the sample time series before training (QN: Quotient Normalisation, "STD": Standard Scaling).**

**A** – Sheep farm| Model trained on healthy 1To1 and unhealthy 2To2

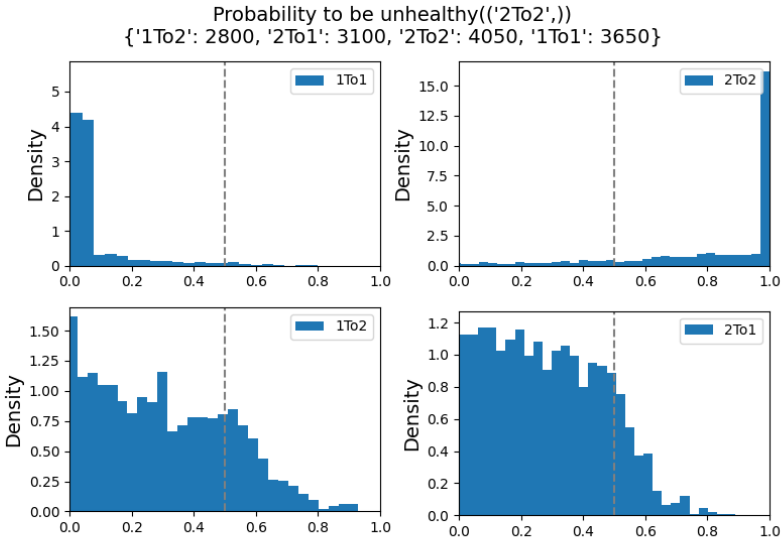


**Fig. S33. Density histograms of prediction results for transitions used in training.** Density histograms are a way to represent the distribution of a set of continuous data. They are similar to regular histograms, which shows the frequency of values within certain intervals (bins) of a continuous variable. However, in a density histogram, the y-axis represents the density of data points within each bin, rather than their frequency. The density is calculated as the number of data points in each bin divided by the total number of data points multiplied by the width of the bin. This ensures that the area under the histogram equals 1, which makes it possible to compare the relative proportions of data in different bins, even if they have different widths.

**A** – Goat farm| Model trained on healthy 1To1 and unhealthy 2To2

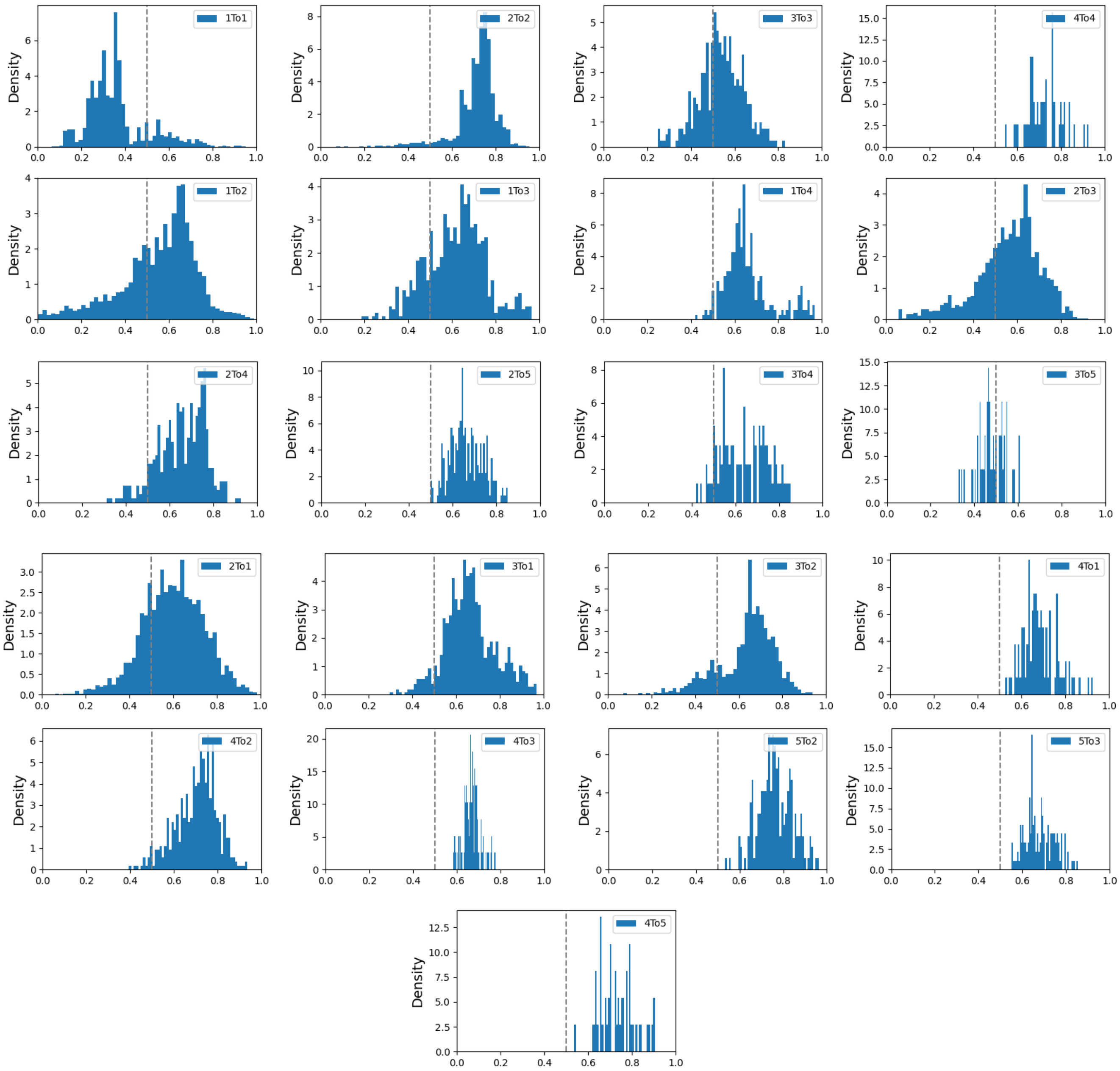


**Fig. S36. SVM predictions on sheep unseen samples.** Density histograms of the model predictions on unseen labels. Each histogram shows the probability output on the x-axis and the density on the y-axis. The dotted grey vertical line marks $Probability = 0.5$. While (A) shows the results for the sheep farm (B) shows the goat. For (B) the FAMACHA transitions tested from top to bottom and right to left are: 1To1, 2To2, 3To3, 4To4, 1To2, 1To3, 1To4, 2To3, 2To4, 2To5, 3To4, 3To5, 2To1, 3To1, 3To2, 4To1, 4To2, 4To3, 5To2, 5To3 and 4To5.

### 6. Regularisation

In the presented regularisation framework, we employed a Support Vector Classifier (SVC) augmented with an exhaustive grid search (GridSearchCV https://scikit-learn.org/stable/modules/generated/sklearn.model_selection.GridSearchCV.html) to optimize hyperparameters, specifically focusing on maximizing the area under the receiver operating characteristic curve (ROC AUC). The hyperparameter space explored includes a logarithmically spaced range for the penalty parameter C (-30 to 10, over 15 points), crucial for regulating the trade-off between margin maximization and misclassification minimization. This exploration is conducted separately for two kernel types: linear and radial basis function (RBF) with a 'scale' gamma parameter (https://scikit-learn.org/stable/modules/generated/sklearn.svm.SVC.html) Utilising the 'scale' parameter for gamma determination, where gamma is set as 1 divided by the product of the number of features and the variance of the input data, proves beneficial in adjusting gamma relative to the dataset's scale and spread.

Setting gamma to $1/(n_features * X.var())$ ensures that in a dataset with many features or high variance, the influence of individual points is moderated, preventing overfitting. For example, in a dataset with 100 features and high variance, this formula reduces gamma, thereby extending the reach of support vectors to form a smoother decision boundary. Conversely, in a dataset with fewer features and lower variance, gamma increases, allowing for a tighter fit around data points.

The regularisation grid search was performed with the next code:

```
svc = SVC(probability=True)
clf = GridSearchCV(svc, parameters, return_train_score=True, cv=5, scoring=’roc_auc’, n_jobs=-1)
parameters = [
    {"C": list(np.logspace(-30, 10, 15)), "kernel": ["linear"]},
    {"C": list(np.logspace(-30, 10, 15)), "gamma": ["scale"], "kernel": ["rbf"]},
]

results = []
for ifold, (train_index, test_index) in enumerate(cross_validation_method.split(X, y)):
    result = classify(train_index, test_index, clf)
    results.append(result)
```

**A** – SVM regularisation C parameter grid search

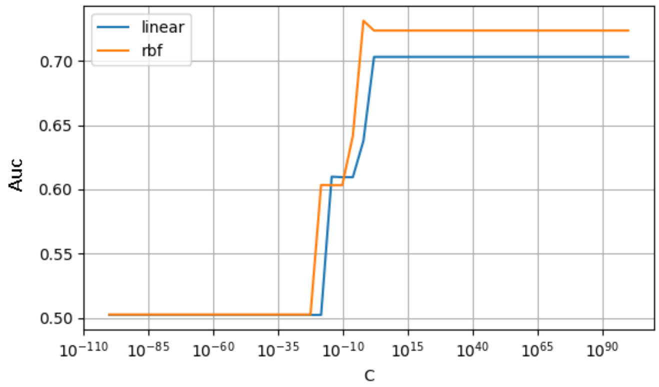


**Fig. S37. SVM regularisation.** Shows the mean auc of all the folds against multiple C values empirically selected for a grid search.

### 7. Interpreting the classifier

**A** –Sheep **B**

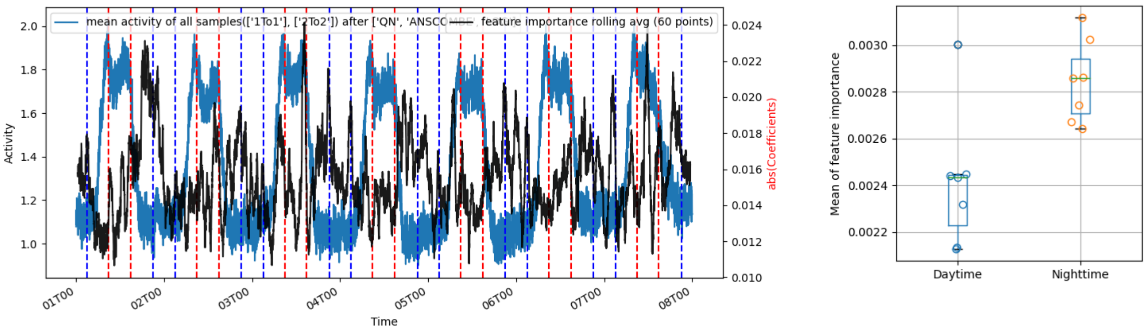


**Fig. S38. Interpretation of the classifiers from Fig. S28.** After fitting the SVM model (A) with the 7 days post-proccessed time series, the blue trace shows the mean of all the samples. We can display the average of the feature importance attributed by the classifier, "black trace". **Statistical analysis** (B) Illustrates the mean importance of all 7 daytime periods vs all 7 night time periods, daytime being defined between the latest sunrise and the earliest sunset, 8am to 4pm, illustrated by the vertical dotted blue and red lines respectively.

**A** –Goats **B**

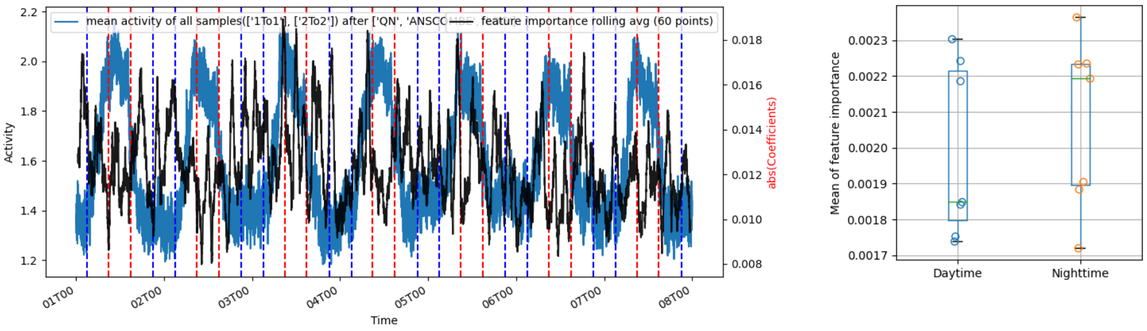


**Fig. S39. Interpretation of SVM classifier** After fitting the SVM model (A) with the 7 days post-proccessed time series, the blue trace shows the mean of all the samples. We can display the average of the feature importance attributed by the classifier, "black trace". **Statistical analysis** (B) Illustrates the mean importance of all 7 daytime periods vs all 7 night time periods, daytime being defined between the latest sunrise and the earliest sunset, 8am to 4pm, illustrated by the vertical dotted blue and red lines respectively.
